# Breakdown of clonal cooperative architecture in multispecies biofilms and the spatial ecology of predation

**DOI:** 10.1101/2022.07.22.501146

**Authors:** Benjamin R. Wucher, James B. Winans, Mennat Elsayed, Daniel E. Kadouri, Carey D. Nadell

## Abstract

Adherence to surfaces and secretion of extracellular matrix, or biofilm formation, is common in the microbial world, but we often do not know how interaction at the cellular spatial scale translates to higher-order biofilm community ecology. Here we explore an especially understudied element of biofilm ecology, namely predation by the bacterium *Bdellovibrio bacteriovorus*. This predator can kill and consume many different Gram-negative bacteria, including *Vibrio cholerae* and *Escherichia coli*. *V. cholerae* can protect itself from predation within highly packed biofilm structures that it creates, whereas *E. coli* biofilms are highly susceptible to *B. bacteriovorus*. Here we explore how predator-prey dynamics change when *V. cholerae* and *E. coli* are growing in biofilms together. We find that in dual species prey biofilms, *E. coli* survival under *B. bacteriovorus* predation increases, whereas *V. cholerae* survival decreases. *E. coli* benefits from predator protection when it becomes embedded within expanding groups of highly packed *V. cholerae*. But we also find that the ordered, highly packed, and clonal biofilm structure of *V. cholerae* can be disrupted if *V. cholerae* cells are directly adjacent to *E. coli* cells at the start of biofilm growth. When this occurs, the two species become entangled, and the resulting disordered cell groups do not block predator entry. Because biofilm cell group structure depends on initial cell distributions at the start of prey biofilm growth, the colonization dynamics have a dramatic impact on the eventual multispecies biofilm architecture, which in turn determines to what extent both species survive exposure to *B. bacteriovorus*.

**Significance Statement:** Bacteria live in multispecies, spatially structured communities ubiquitously in the natural world. These communities, or biofilms, have a strong impact on microbial ecology, but we often do not know how cellular scale interactions determine overall biofilm structure and community dynamics. Here we explore this problem in the context of predator-prey interaction, with two prey species – *Vibrio cholerae* and *Escherichia coli* – being attacked by the bacterial predator *Bdellovibrio bacteriovorus*. We find that when *V. cholerae* and *E. coli* grow together in biofilms, the architectures that they both produce change in ways that cannot be predicted from looking at each prey species alone, and that these changes in cell group structure impact the community dynamics of predator-prey interaction in biofilms.

## Introduction

Most organisms do not naturally live in isolated monocultures but rather in communities composed of many species, and microbes are no exception (1–4). Bacterial communities are present ubiquitously including sinking detritus particles in aquatic environments, deep-sea hydrothermal vents, rhizosphere habitats, animal digestive tracts, plant roots and shoots, fouling surfaces across human industry, and many types of chronic infections (5–19). In many of these contexts, surface attachment and growth in large cell groups, or biofilm formation, are important strategies for sequestering limited space and nutrients, as well as protecting against common biotic and abiotic threats (20–25). Biofilms are small scale ecosystems encased in a wide range of secreted polymeric substances that control cell-cell and cell-surface engagement (26–32). The benefits of biofilm formation have been examined in many contexts, showing their capacity for public goods sequestration, exclusion of newly arriving competitors, predation protection, and antibiotic tolerance (33–37). The precise costs and benefits of biofilm formation vary across species, as do the multicellular architectures that emerge from the combination of cell growth, matrix secretion, and environmental feedbacks (23, 27, 29, 38–46). In what few cases have been examined, multispecies biofilms create structures that may be distinct from those found in single species biofilms of the community constituents in isolation (47–49). Understanding the connections between biofilm architecture and microbial community ecology at different scales remains an important area for ongoing work on numerous topics, including crossfeeding relationships, diffusible and contact-mediated toxin antagonism, resilience against antibiotics in therapeutic contexts, industrial and medical surface degradation, and others (50–56).

Predation within biofilms is a broad sub-class of microbial ecology that has received relatively little attention with high resolution imaging and analysis. While many predators, such as phages, can usually attack only a small number of prey species, the ubiquitous predator *Bdellovibrio bacteriovorus* can be considered a generalist (36, 57–60). It is not known to rely on a specific receptor for cell entry and can prey on a variety of proteobacteria (8, 59, 61–63). However, the extent of its predation on biofilm-dwelling target cells appears to vary widely between prey species, and the mechanisms underlying this variability remain mostly unknown.

Studies using macroscopic measurement techniques have reported the susceptibility of several biofilm-producing species to *B. bacteriovorus* predation (57). For example, *Escherichia coli* and *Pseudomonas fluorescens* can be largely or entirely consumed by *B. bacteriovorus* in laboratory biofilm culture (36). We recently documented a different outcome in *Vibrio cholerae* biofilms, which can protect themselves from predator exposure via their highly packed cell arrangements that occur after the prey cell groups grow beyond several hundred cells (35). Given that we have examples of prey species whose biofilm structure protects them from *B. bacteriovorus* predation, and other prey whose biofilms confer little or no protection at all, we were curious as to what would happen to predator-prey dynamics in multi-species prey biofilm contexts. How often and to what extent does the cellular architecture of the two species depart from what is normally observed in monoculture? How do any of these changes influence the susceptibility of different prey species to *Bdellovibrio* predation, and overall predator-prey population dynamics?

We chose two proteobacteria – *V. cholerae* and *E. coli* – to study these questions. Both species have been isolated from biofilms in the same environments in proximity to humans, and they have been documented to co-infect hosts (64–69). Their respective biofilm formation mechanics have also been very well characterized. *V. cholerae* forms dense, tightly packed cell groups with radially aligned cell arrangements, whose structure depends on the matrix proteins RbmA, RbmC, and Bap1, as well as VPS, its primary known matrix polysaccharide (27, 28, 40, 70–72). This highly packed cell group architecture is critical for protection from phages and from *B. bacteriovorus* (35, 37). The cell packing required for predator protection does not occur immediately as biofilm growth begins, though, leaving nascent *V. cholerae* cell groups open to predation. *E. coli* in contrast forms its biofilms with many different matrix components including cellulose, polyglycolic acid, colanic acid, Type 1 fimbriae, flagellar filaments, and curli fiber proteins (73). *E. coli* matrix architecture, and curli protein in particular, have been shown to confer protection against phage exposure, but other prior work has indicated even structurally mature *E. coli* biofilms are not protected from *B. bacteriovorus* (57, 74). Here we use single cell resolution microscopy to examine the structure of these two prey species in monoculture and dual-culture biofilms, finding that the multispecies context causes unexpected changes in biofilm architecture that in turn alter predator-prey interaction and the overall population dynamics.

## Results

### Predation in dual-species prey biofilms has opposite fitness effects for *E. coli* and *V. cholerae*

We engineered *V. cholerae* N16961, *E. coli* AR3110, and *B. bacteriovorus* 109J to constitutively produce the fluorescent proteins mKate2, mKO-κ, and GFP, respectively, so that they could be distinguished for live confocal microscopy. N16961 is naturally repressed for Type 6 Secretion System activity in most conditions, and so does not kill *E. coli* via this mechanism in our experiments (75–77). Overnight cultures of the two prey species were both normalized to OD_600_=1.0 before inoculating them at a 1:1 ratio into poly-dimethylxilosane microfluidic flow devices bonded to coverslip glass (see Methods). In parallel we performed mono-culture experiments in which *V. cholerae* and *E. coli* were introduced to chambers on their own. Cells were allowed to colonize the underlying glass surface in stationary conditions for 1 h, after which M9 minimal media with 0.5% glucose was introduced into the chambers at 0.2 μL/min (average flow velocity = 90 μm/s). After 48 h of growth in co-culture, groups with varying composition of both species could be found distributed throughout the chambers (SI Figure S1A), and prior to exposure to *B. bacteriovorus*, the two prey species equilibrated at frequencies of ~90% *V. cholerae* and ~10% *E. coli* (Figure 1; SI Figure S1A). Following 48 h of prey biofilm growth, we introduced *B. bacteriovorus* under continuous flow for 1h, followed by a return to influx of sterile M9 media for both the dual culture chambers and the monoculture controls. When exposed to *B. bacteriovorus* in a mono-species context, *V. cholerae* survives within cell groups that have reached high cell packing, as we have shown previously (35). 48 h following predator exposure, monoculture *V. cholerae* biofilms maintain net positive growth (Figure 1A, B). By contrast, and consistent with prior reports, *E. coli* biofilms in monoculture exhibit little to no survival in the presence of *B. bacteriovorus*, with viable prey biomass (that is, *E. coli* cells without *B. bacteriovorus* inside or attached to them) falling nearly to zero 48 h after predator introduction (Figure 1A, B; SI Figure S1B) (36).

**Figure 1.**
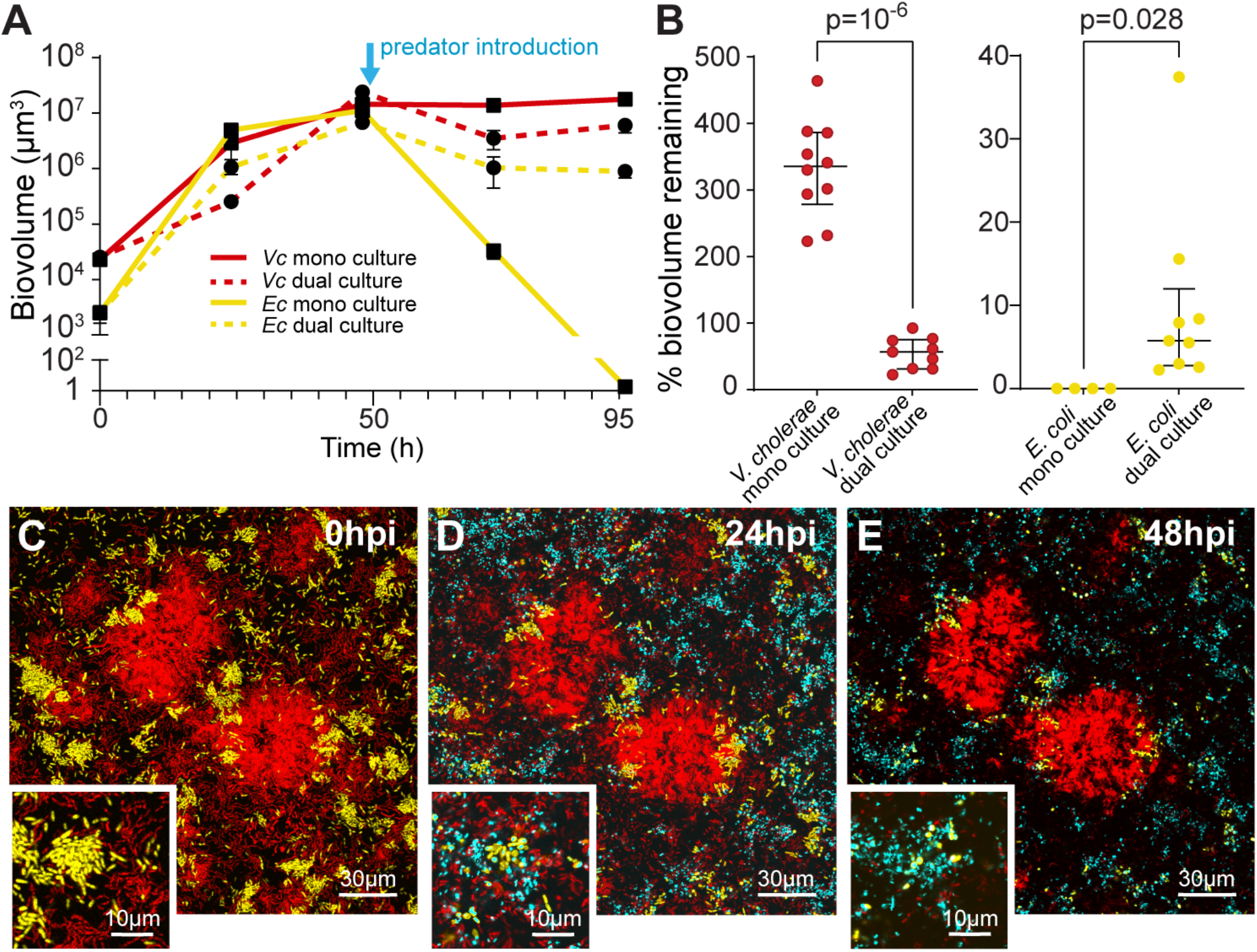
Population dynamics in monoculture and dual culture biofilms of *V. cholerae* (red) and *E. coli* (yellow) undergoing predation by *B. bacteriovorus* (cyan). Biofilms of *V. cholerae* and *E. coli* were grown for 48 h prior to *B. bacteriovorus* predator exposure. (A) Population dynamics of each prey species in monoculture and dual culture biofilm growth (*n* = 4). (B) Percent change in prey biovolume, a direct proxy for population size, 48 h after predator introduction relative to just prior to predator introduction, both in single species prey biofilm controls and dual species prey co-culture biofilm conditions. (*V. cholerae* monoculture *n* = 10; *V. cholerae* dual culture *n* = 9; *E. coli* monoculture *n* = 4; *E. coli* dual culture *n* = 9; plots denote medians with interquartile ranges). Pairwise comparisons were performed with Mann-Whitney U-tests. (C-E) Representative images of dual culture biofilms at (C) 48h after initial prey inoculation (just prior to predator introduction; ‘hpi’ denotes ‘hours post introduction’ of predators), (D) 24h post introduction of predators, and (E) 48 h post introduction of predators. Inset frames show regions with details of predators entering host cells and forming rounded bdelloplasts, indicating active predation. Images are single optical sections just above the glass substratum, showing the bottom layers of the biofilms.

If *B. bacteriovorus* predates *V. cholerae* and *E. coli* independently of one another in dual prey species biofilms, we would expect residual *V. cholerae* biomass and elimination of all *E. coli* subpopulations, as seen in the prey monoculture experiments above. However, examining the population dynamics quantitatively, we found a significant increase in *E. coli* survival and a significant decrease in *V. cholerae* survival following predator introduction in the dual-species condition relative to the single species controls (Figure 1). These results imply nonlinear interactions between *V. cholerae*, *E. coli*, and *B. bacteriovorus*, which alter prey population dynamics in a manner that the prey mono-species controls cannot predict. Below we explore why *E. coli* fares better in dual-species biofilms against predator exposure, and why *V. cholerae* fares worse, relative to their respective single species prey conditions.

### *E. coli* gains protection from predator exposure while embedded within *V. cholerae* cell groups

Prior to introduction *of B. bacteriovorus* predators, we noticed that the spatial distributions of *V. cholerae* and *E. coli* in dual-culture prey biofilms were heterogeneous on scales of 10-100 μm, and in some locations, there were groups of *E. coli* cells that had been enveloped by along the basal layers of expanding colonies of highly packed *V. cholerae*. 48 h following introduction of predators, virtually all surviving *E. coli* were those embedded along the bottom of packed *V. cholerae* biofilms in this manner (Figure 2A; SI Figure S2; see also SI Figure S4D). As previously documented, *V. cholerae* biofilm clusters reach a cell packing threshold that blocks predator entry and allows the interior cells to survive (35). *E. coli* enveloped within *V. cholerae* groups that have reached this threshold appear to gain the predation protection conferred by *V. cholerae* biofilm structure. To assess this point in more detail we quantified the extent of *E. coli* predation and *V. cholerae* fluorescence in proximity to *E. coli* in these images; *E. coli* with high *V. cholerae* fluorescence in close proximity clearly also experienced the least predation (Figure 2B,C; this result is clarified with further quantitative detail in the next section). Additionally, protection of *E. coli* within *V. cholerae* cell groups was dependent on the high-density structure that wild type *V. cholerae* creates. When the experiment was repeated with a mutant strain of *V. cholerae* that cannot produce RbmA – a matrix component required for the tight cell packing found in mature biofilms – *B. bacteriovorus* could freely enter and access both *V. cholerae* and *E. coli* (SI Figure S3A,B). In this case, almost all cells of both species were killed off by the predators (SI Figure S3C,D). In a parallel study illustrating the generality of these results, we show that *E. coli* embedded along the bottom layers of expanding *V. cholerae* biofilm clusters are also protected from exposure to the obligate lytic phage T7 and temperate phage l in a manner dependent on the high packing structure of *V. cholerae* biofilms (78).

**Figure 2.**
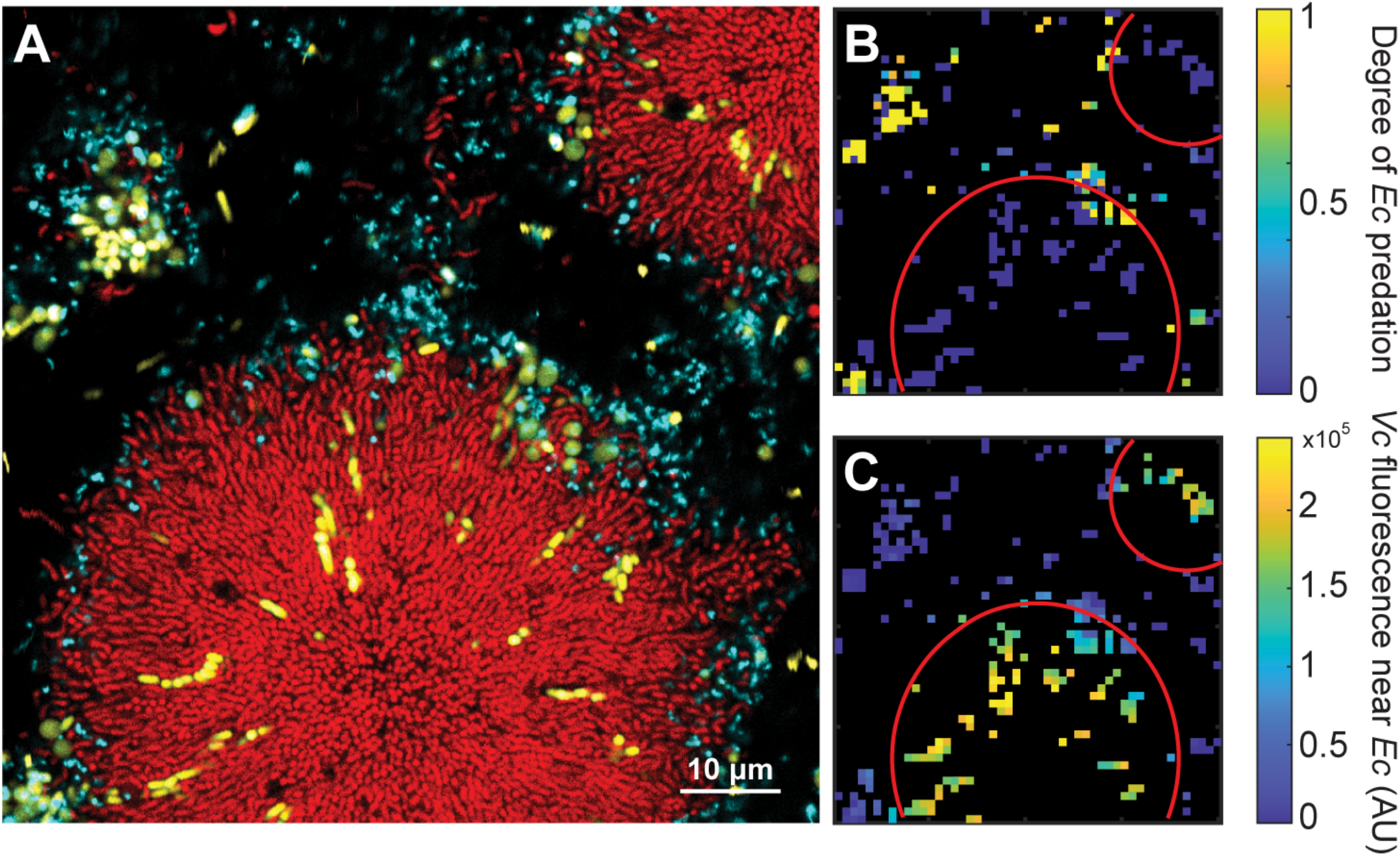
*E. coli* (yellow) enveloped within highly packed *V. cholerae* biofilms (red) can be protected from *B. bacteriovorus* (cyan) exposure. (A) Representative image demonstrating the ability of highly packed *V. cholerae* biofilms to protect *E. coli* biomass from access by *B. bacteriovorus*. The image is a single optical section just above the glass substratum. (B) Heatmap of the degree of predation on *E. coli*, quantifying raw data from panel A. Red circles denote boundaries of highly packed *V. cholerae* cell groups. (C) Heatmap of *V. cholerae* fluorescence within 5μm of each unit of segmented biovolume of *E. coli* from panel A.

Although *E. coli* cells embedded within *V. cholerae* biofilms gain predator protection, it was not clear if they remain viable, or if they are able to subsequently disperse to colonize downstream locations. If not, then the protection *E. coli* gains from *B. bacteriovorus* exposure within *V. cholerae* cell groups would not necessarily translate to meaningful fitness gains on longer time scales. We explored this question by inducing a disturbance regimen following *B. bacteriovorus* predation of co-culture prey biofilms, allowing the *V. cholerae* and *E. coli* to colonize new microfluidic devices from the effluent exiting the initial chambers. We could show that almost all *E. coli* outside the periphery of highly packed *V. cholerae* colonies were killed by *B. bacteriovorus* exposure, and that the remaining *E. coli* that were protected while embedded within *V. cholerae* colonies remained viable. When these surviving *E. coli* were dislodged by disturbance, they could successfully colonize new locations downstream (SI Figure S4).

The predation protection gained by *E. coli* cell groups embedded within highly packed *V. cholerae* biofilm clusters explains the increase in *E. coli* survivorship that we originally observed in dual culture. We note however that the *V. cholerae* cells within these cell groups are also protected from *B. bacteriovorus* predation (Figure 2), so the results so far do not yet explain why *V. cholerae* survivorship declines in coculture with *E. coli* under predation pressure.

### Co-culture with *E. coli* can lead to breakdown of *V. cholerae* biofilm architecture

In the previous section we made note of highly packed *V. cholerae* biofilms into which *E. coli* had become embedded and gained protection from *B. bacteriovorus* exposure. These colonies appear to behave in much the same way as mono-species *V. cholerae* biofilms, as they contain large continuous groups of *V. cholerae* in their ordered radial alignment and tight packing, albeit with pockets of *E. coli* along the glass substratum included as well (Figure 3A, B). This growth pattern was not the only kind that emerged in prey co-culture experiments, however: there was a second, qualitatively distinct colony architecture in which *V. cholerae* and *E. coli* were homogenously mixed together (Figure 3A,C). These colonies were disordered in comparison with the radial alignment in ordered *V. cholerae* cell groups, with visibly reduced cell packing density. *B. bacteriovorus* could enter throughout these disordered cell groups, gaining access to and killing most or all *V. cholerae* and *E. coli* cells within them (Figure 3A,C).

**Figure 3.**
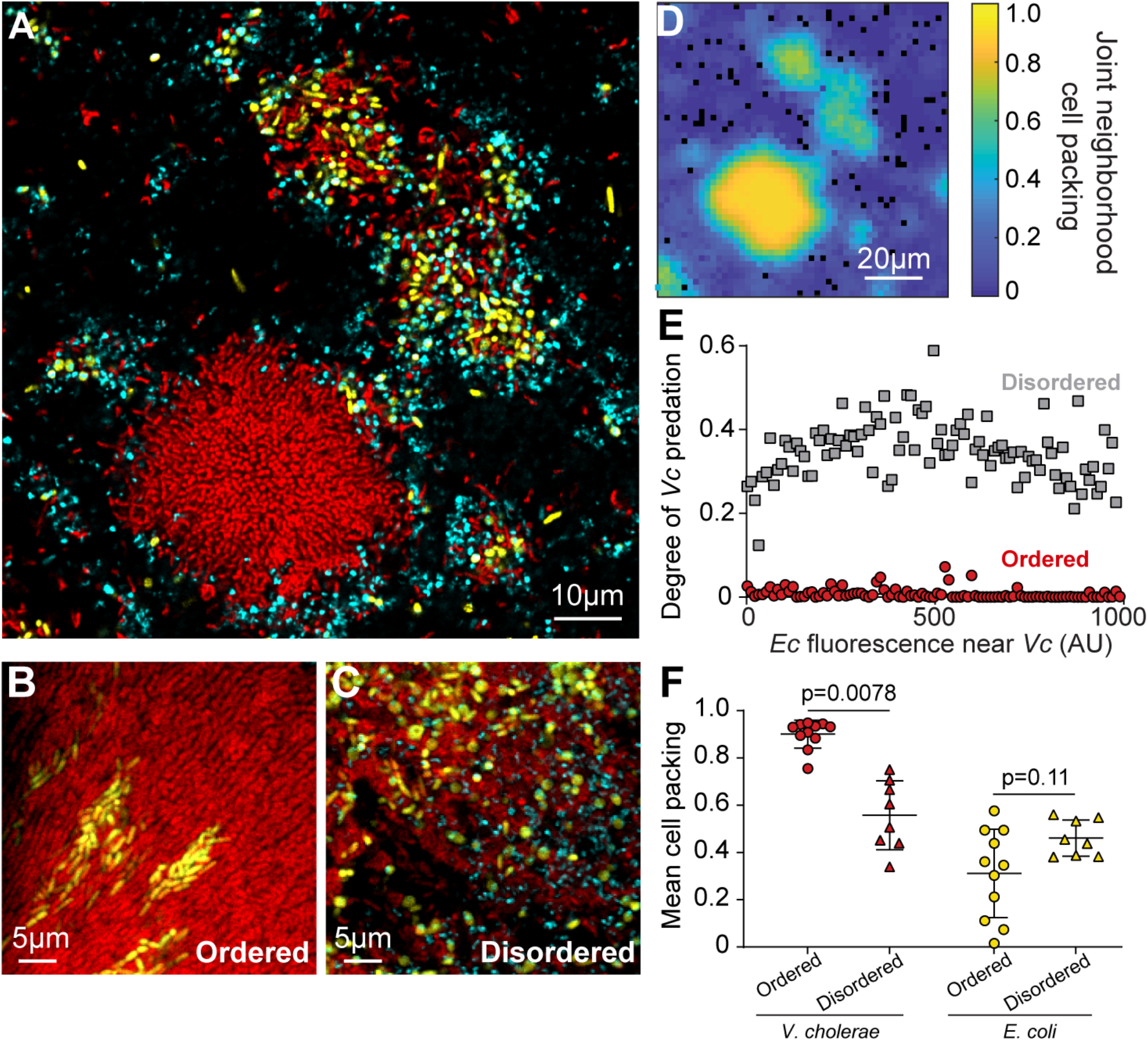
*V. cholerae* (red) and *E. coli* (yellow) exhibit two distinct joint biofilm morphologies in co-culture that strongly affect *B. bacteriovorus* (cyan) predation susceptibility. (A) Representative image of both biofilm dual species cell group types, which can occur in close proximity. The structure we term ‘ordered’ more closely resembles the architecture *V. cholerae* produces on its own and is shown on the lower left in this image. The novel, well-mixed structure, which we term ‘disordered’, is shown on the upper right. (B-C) Additional higher magnification images detailing the architecture of ordered and disordered colony morphologies. (D) Heatmap of the combined two species neighborhood cell packing values for the image in panel A. (E) Scatterplot of the degree of predation on *V. cholerae* as a function of the fluorescence of *E.coli* in proximity to *V. cholerae* biomass. The data are split according to whether they are from ordered architecture colonies (*n* = 11), or disordered colonies (*n* = 9). (F) Within-species cell packing for *V. cholerae* (red) and *E. coli* (yellow) in 48 h incubated cell groups of each structure type (ordered *n* = 11; disordered *n* = 9; plots denote medians with interquartile ranges). Pairwise comparisons were performed by Mann-Whitney U-test.

To study the differences between the ordered and disordered colony structure in more detail, we isolated examples of each for further analysis (Figure 3B,C). The relative abundance of *V. cholerae* looked to be somewhat lower in disordered colonies (Figure 3B,C, see also next section for quantitative detail). The combined cell packing of the two species together was indeed diminished in disordered colonies (Figure 3D), falling below the threshold necessary for predation protection in ordered *V. cholerae* colonies (35). An alternative but not mutually exclusive factor for predation susceptibility for *V. cholerae* could be abundance of *E. coli* in their immediate vicinity. However, measurements of the relationship between *V. cholerae* predation and local abundance of *E. coli* in the two colony types clearly shows that overall colony structure is the dominant factor controlling the extent of predation by *B. bacteriovorus* (Figure 3E). A similar reciprocal analysis of *E. coli* predation as a function of colony type and local abundance of *V. cholerae* gave the same outcome (Figure S5). The shift in architecture between ordered and disordered cell groups also appeared to be driven entirely by the loss of *V. cholerae*’s ability to produce highly packed clusters, as it normally does on its own. Examining the cell packing of groups formed after 48 h supports this interpretation, as *V. cholerae* within-species cell packing with respect to itself (in contrast with joint cell packing of the two species together noted in (Figure 3D) shifted between ordered versus disordered biofilm types (Figure 3F). *E. coli* within-species cell packing, on the other hand, was not statistically different in ordered versus disordered cell groups (Figure 3F). On longer time scales, it should be noted, *E. coli* cell groups that have been enveloped within ordered *V. cholerae* colonies can also be driven to high within-species cell packing by the confinement imposed by *V. cholerae* biofilm architecture (78).

Our results thus far indicate that in mixed prey biofilms of *V. cholerae* and *E. coli*, a fraction of the *E. coli* along the basal surface becomes engulfed within *V. cholerae* cell groups with ordered high packing structure that protects both species from *B. bacteriovorus*. This observation explains the improvement in *E. coli* predation survival in dual culture relative to monoculture biofilms. On the other hand, a fraction of the *V. cholerae* population becomes entangled with *E. coli* in well-mixed colonies that fail to develop *V. cholerae*’s normal packing structure, instead growing into disordered, loosely assembled groups that are fully susceptible to predation. This observation clarifies why *V. cholerae* predation protection declines in biofilm co-culture with *E. coli*.

### Mixed species cell groups follow distinct trajectories depending on surface colonization conditions

To better understand how the highly packed versus disordered dual-species colonies originate, we ran new experiments tracking biofilm growth at 1 h intervals from the initial stages of surface colonization to clusters containing hundreds of cells at 48 h. Examples of the two cell group types were found by visual inspection at 48 h and then traced back to their initial conditions corresponding to 9 h after the start of incubation (Figure 4A,C). On larger spatial scales than shown in Figure 4, both types of colony morphology could be found in close proximity (Figure 3A, SI Figure S6). The two colony types could be reliably distinguished by their combined cell packing, which was systematically lower in the core regions of disordered colonies (Figure 4B,D, see below for temporally resolved detail). The two colony structures were also consistently different in total biovolume and species composition at 48 h, with ordered high-packing cell groups growing to larger population sizes and containing higher relative abundance of *V. cholerae* compared to *E. coli* (Figure 4 E,F).

**Figure 4.**
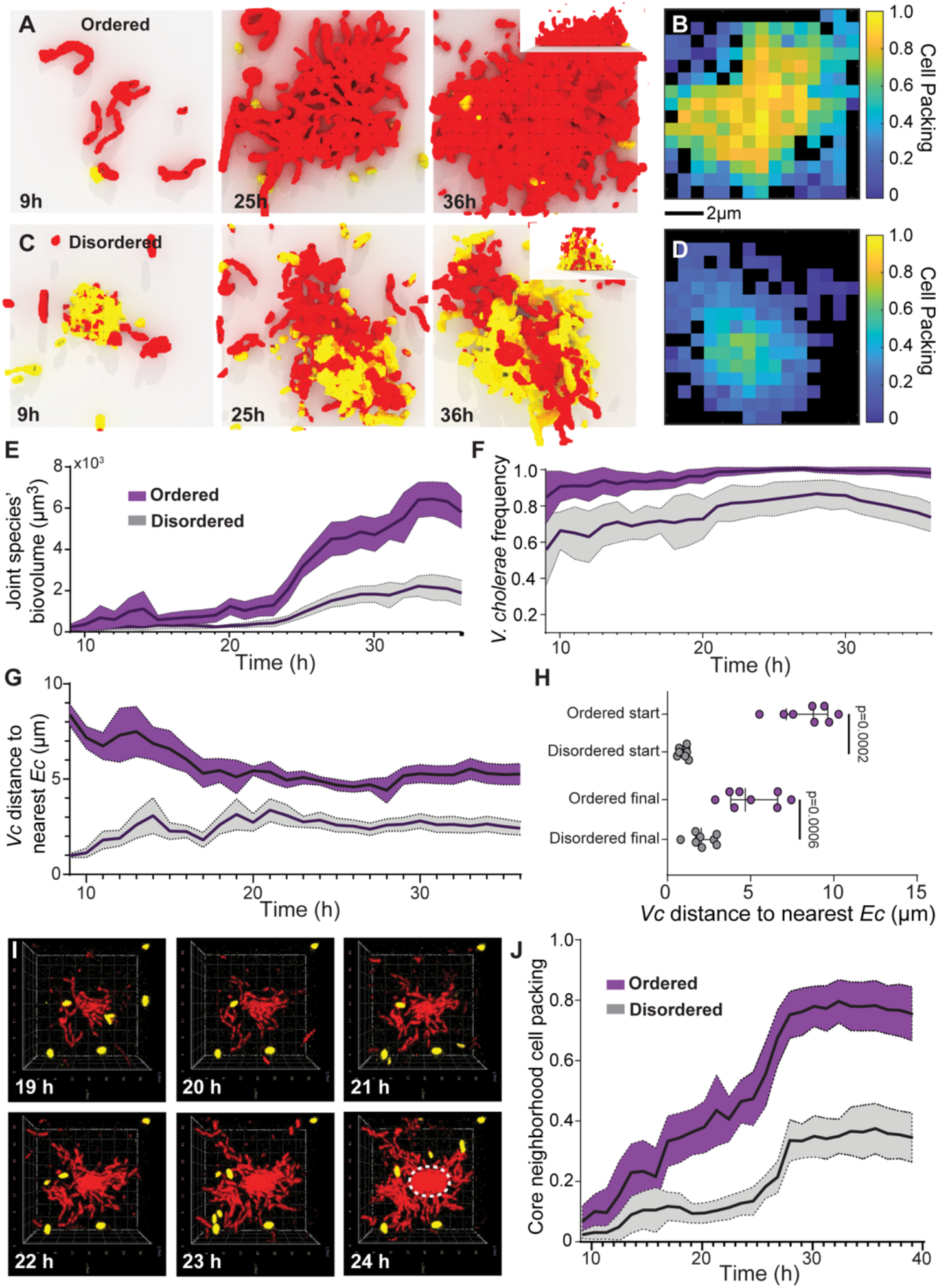
The initial distance between *V. cholerae* (red) and *E. coli* (yellow) cells best distinguishes colonies that will become highly packed and predation-protected versus disordered and predation-susceptible. (A) Time lapse of 3D renderings of an example ordered cell group with (B) a heatmap for neighborhood cell packing at the 36 h time point. (C) Time lapse of 3D renderings of an example disordered cell group with (D) a heatmap for neighborhood cell packing at the 36 h time point. Renderings in panels A and C are 31μm × 31μm × 14μm) (LxWxD). (E-G) Time courses for (E) combined biovolume of both species, (F) *V. cholerae* frequency, (G) Average distance of *V. cholerae* cells to nearest *E. coli* (*n* = 8 for each colony type). (H) Statistical comparison of *V. cholerae* distance to nearest *E. coli* between biofilm types at the start and end of courses in G (Mann-Whitney U tests with *n* =8; plots denote medians with interquartile ranges). (I) Top-down view of a *V. cholerae* cluster as a core high-packing region nucleates (each rendering is 31μm × 31μm - 14μm) (LxWxD). The secondary biofilm front that forms the boundary of this core high packing region is denoted with a dotted white circle in the 24 h time point image. (J) Time courses of neighborhood cell packing in the core regions (within 10μm of colony center) of both colony types.

Though the two biofilm architectures could be quantitatively separated in several respects after they had grown to several hundred cells in size, the mean distance between *V. cholerae* and *E. coli* cells was the only factor we could discern at early time points that differentiated cell groups that would later become ordered, highly packed biofilms versus disordered cell groups (Figure 4G; SI Figure S7). The two cell group types in fact begin and remain on different trajectories with respect to *V. cholerae*-*E. coli* distance across all replicate colony time series acquisitions. *V. cholerae* that produced ordered, highly packed architecture were on average 8 μm away from the nearest *E. coli* cell at early time points, while those that became disordered clusters were only 1 μm from the nearest *E. coli* cell at early time points (Figure 4G,H). This observation suggests that the colonization conditions at the beginning of biofilm growth, as surface-attached cells are just starting to divide in place, are crucial for the eventual consolidation of *V. cholerae* packing architecture.

Reviewing the data obtained from time series of the two dual species colony structures, we also noted an important transition that occurs in ordered high-packing *V. cholerae* cell groups. In our culture conditions at ~19-26 h, a core region of highly packed *V. cholerae* cells is nucleated (Figure 4I). This process creates a secondary front of structural consolidation that lags behind the outermost, less densely packed growth front that represents the interface between the growing colony and the surrounding liquid medium (Figure 4B). This secondary front bounding the central core biofilm region has been observed previously (28, 29, 79), and it corresponds to the portion of the cell group with sufficiently high cell density to provide predator protection (35). In disordered colonies containing well-mixed *V. cholerae* and *E. coli*, the nucleation of this high-density core never occurs. This difference between the two cell group types can be tracked quantitatively via the cell packing of the inner core of each (Figure 4J). In ordered colonies, once the highly packed core was initiated, it was stable over time, and interruption of this core nucleation process only occurred if *V. cholerae* and *E. coli* cells happened to begin growing in close proximity from the start of biofilm formation. Allowing *V. cholerae* to grow on its own for 48 h, followed by invasion of *E. coli* into the biofilm environment, never led to any observable disruption of highly packed *V. cholerae* groups. Introduced planktonic *E. coli* cells could not invade *V. cholerae* biofilms and were completely susceptible to predation if *B. bacteriovorus* was later added to the system (SI Figure S8).

Our results here highlight critical points for the production of *V. cholerae* biofilm structure with its characteristic packing and cell alignment architecture (27–29, 34, 79), which in turn are necessary for protection from *B. bacteriovorus*(35). If *V. cholerae* cells are sufficiently isolated during early biofilm formation, they can grow, divide, and secrete biofilm matrix components that effectively coordinate their normal architecture. However, if *V. cholerae* cells are too close to *E. coli* at the start of biofilm formation, the two species become entangled in the process of growth and division in a manner that interrupts the longer-term production of cell group architecture that *V. cholerae* normally produces on its own. The disruption of the structure that *V. cholerae* typically produces suggests that if *E. coli* is in close enough proximity at early stages of growth, *V. cholerae* either does not produce biofilm matrix normally, or it does produce matrix, but with a disruption of the cell-cell orientation and matrix localization that *V. cholerae* would obtain on its own.

To begin parsing these possibilities, we performed additional biofilm co-culture experiments in which *V. cholerae* produced FLAG-labeled versions of its primary matrix proteins RbmA, RbmC, and Bap1, which confirmed that *V. cholerae* is indeed producing biofilm matrix in direct proximity to *E. coli* (SI Figure S9A-C). We also found, surprisingly, that RbmC localizes substantially above background around *E. coli* cells (SI Figure S9B,D). This result was also supported by experiments in which cell free *V. cholerae* supernatant containing RbmC-FLAG was added to *E. coli* biofilms growing in monoculture (SI Figure S10). RbmC is a diffusible matrix protein that participates in the internal architecture of *V. cholerae* clonal cell groups but also, importantly, contributes to binding of the cell groups to the underlying surface (27, 29, 71). The mechanical details of binding between the group and surface are important for the transition from lateral expansion of flat monolayers to the extension of *V. cholerae* biofilms into 3-D space and subsequent packing architecture (29, 80). Accumulation of RbmC around *E. coli* does not appear to diminish the amount of RbmC that *V. cholerae* itself accumulates relative to monoculture conditions (SI Figure S9E). Nevertheless, the localization of this matrix protein around *E. coli* in close proximity to *V. cholerae* may potentially contribute to the disruption of normal *V. cholerae* cell group architecture that we observe when the two species begin biofilm growth directly adjacent to each other. The precise physical and biochemical details of how multispecies biofilm architecture quantitatively and qualitatively departs from clonal biofilm architecture will be an important area for future work.

### Surface colonization strongly impacts population dynamics via its influence on biofilm architecture

Our results above suggest that the average distance between *V. cholerae* and *E. coli* cells at the start of biofilm growth should directly determine the relative occurrence of ordered, highly packed *V. cholerae* groups that envelope pockets of surface-attached *E. coli* as they expand, as opposed to disordered dualspecies cell groups. As highly packed *V. cholerae* cell groups are protected from *B. bacteriovorus*, while disordered colonies are not, our observations lead to the ecological prediction that initial surface colonization conditions, via their impact on the relative proportion of highly packed versus disordered dual species colonies, can substantially change the population dynamics of predator-prey interaction. In other words, the initial surface coverage should indirectly determine the overall impact of predation on survival of both species via its direct impact on colony structure development. We tested this prediction by inoculating two sets of two-species chambers with relatively low or high surface colonization density (20% versus 60% surface coverage, respectively, SI Figure S11A). Low or high initial density alters the distributions of distance between *V. cholerae* and *E. coli* cells (SI Figure S11B), in turn leading to large amounts of highly packed *V. cholerae* colonies containing small numbers of *E. coli* (low initial density), or almost exclusively disordered, mixed colonies containing homogeneously distributed *V. cholerae* and *E. coli* (high initial density) (Figure 5A,B).

**Figure 5.**
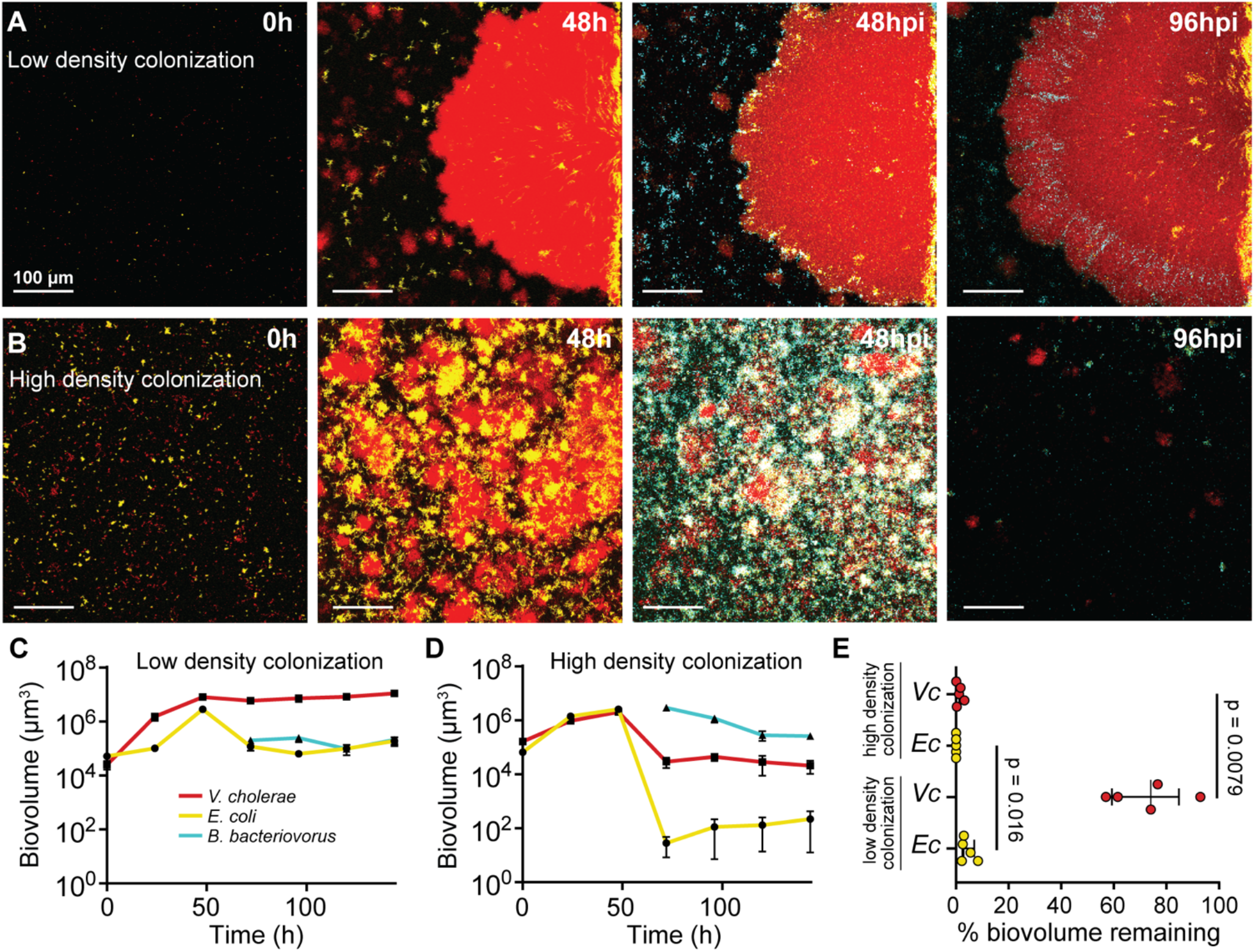
Initial surface colonization density alters the fraction of highly packed versus disordered colonies of *V. cholerae* (red) and *E. coli* (yellow), which in turn alters overall population dynamics and *B. bacteriovorus* (cyan) predator survival for both prey. (A,B) Representative image sets of the low density (A) and high density (B) colonization conditions at initial surface colonization through 96 hours post predator introduction. (C,D) Biovolume of all three species as a proxy for their population dynamics in the (C) low-density initial condition and (D) high-density initial condition (*n* = 5; plots denote medians with interquartile ranges). (E) Per cent of each prey species remaining at the end of the predation experiments in the low and high colonization density initial conditions (*n* = 5; plots denote medians with interquartile ranges). The percentage remaining is calculated as the biovolume of each prey species at the last time point (146 h) relative to the biovolume that was present just prior to the introduction of *B. bacteriovorus* predators. Pairwise comparisons are Mann-Whitney U tests.

As anticipated, low versus high initial surface occupation led to distinct survival outcomes for both prey species after the introduction of *B. bacteriovorus*. In the low initial density condition, the majority of *V. cholerae*, as well as the smaller groups of *E. coli* embedded in their packed biofilms, survive predator exposure (Figure 5C,E,F). By contrast, in the high initial density conditions predominated by disordered groups of *V. cholerae* and *E. coli*, almost the entire dual species prey community is killed off following the introduction *B. bacteriovorus* (Figure 5D-F). Taken together, these data are consistent with our prediction that surface colonization conditions determine the relative amounts of ordered versus disordered biofilm cell groups of *E. coli* and *V. cholerae*, which in turn governs the survival of both prey species under *B. bacteriovorus* predation.

## Discussion

Exploration of multi-species biofilm communities using live high-resolution imaging is crucial to understanding microbial ecology at the spatial scale on which cell-cell interactions occur (11, 20, 23, 81–90). Here we tracked the spatial population dynamics of the bacterial predator *B. bacteriovorus* in dual-species prey biofilms of *V. cholerae* and *E. coli*, finding that the survival rates of both prey species are substantially altered, but in opposite directions, when they are growing together. *V. cholerae* produces biofilm cell clusters that reach a cell packing density threshold past which *B. bacteriovorus* cannot enter, protecting the prey within. *E. coli* can become enveloped along the basal layers of these highly packed structures, co-opting predator protection from *V. cholerae* and increasing *E. coli* survival relative to when growing on its own. By contrast, in dual species biofilms, a fraction of *V. cholerae* becomes entangled with *E. coli* early during biofilm growth, leading to an alternate structure that is homogeneously mixed, disordered, and loosely packed. These disordered cell groups do not block predator cell entry, and all prey within them are killed by *B. bacteriovorus*. As a result of these biofilm structural dynamics, *V. cholerae* survival decreases in co-culture with *E. coli* relative to when growing on its own. At any given location, which of these two alternative cell group structures emerges depends on the initial distance between *V. cholerae* and *E. coli* cells that have attached to the underlying surface. Surface colonization patterns therefore determine the relative occurrence of predation-protected cell groups versus susceptible cell groups, and the overall rates of *B. bacteriovorus* predator survival for each prey species.

This study makes explicit that the cellular arrangement and tightly packed structure of clonal *V. cholerae* groups can operate as a type of public good (39, 91) that confers predator protection to the cells within (among many other benefits (30, 30, 33–35, 55, 74, 92–96)). Other species – here, *E. coli*, whose mono-species biofilms are susceptible to *B. bacteriovorus* – can take advantage of this protective architecture when small groups of them become enveloped by expanding, highly packed biofilms of *V. cholerae*. By contrast, if too many *E. coli* cells are present in close enough proximity to *V. cholerae* at the start of biofilm growth, then *V. cholerae* cannot initiate its normal cell group structure, and the public good benefit of predation protection completely breaks down in that location. It is notable that the spatial architecture of biofilm-producing bacteria can manifest as a public good that is exploitable across species in this manner. In this case the stability of *V. cholerae* cooperative architecture depends on the initial surface population density, which determines whether *V. cholerae* cell lineages have enough space to nucleate the highly packed core regions of expanding biofilm clusters before encountering cells of other species. Though distinct in mechanistic detail, this example should fall under related social evolution principles as other kinds of microbial cooperation that provide benefits in a distance-dependent manner. Recent work has highlighted in detail how the population dynamics and evolutionary stability of this class of cooperative behavior depends on the spatial range of cooperative sharing, the population/community composition, and spatial cell arrangements during early biofilm growth (33, 34, 39, 92, 95, 97).

The interplay of *V. cholerae*, *E. coli*, and *B. bacteriovorus* in co-culture emphasizes that the population dynamics of different species in a community can depend quite strongly on the cellular resolution details of biofilm structure, which in turn can differ in unexpected ways between mono-species and multi-species systems. In recent years microbiologists have made tremendous strides in understanding the cellular and molecular nuances of biofilm architecture and their relationship to microbial ecology and evolution. By necessity for tractability in many cases, much of this work has focused on one species at a time. Our experiments here highlight how new and interesting questions about the drivers of biofilm structure, and the relationship between biofilm structure and community ecology, can arise from modest increases in complexity with multi-species systems. Here, it appears as though *E. coli* – if adjacent to *V. cholerae* at the start of biofilm growth – may interfere with normal localization of at least one component of the *V. cholerae* matrix, which could then contribute to the qualitative differences in ordered versus disordered architectures that appear later during biofilm growth.

The connection between initial surface coverage and multispecies prey biofilm architecture, and the additional connection between biofilm architecture and predator exposure, together lead to an interesting dependence between early biofilm growth conditions and predator-prey ecology. It would be fruitful to explore how and when these relationships generalize to other species combinations and biofilm environmental growth conditions. Where prior studies have analyzed multispecies biofilms at high resolution, they have also indicated important consequences for community structure and environmental impacts (11, 12, 22, 47, 48, 98–102). A notable recent example examined the detailed structure of multispecies biofilm communities growing as plaque associated with dental caries (48). Kim et al. showed that *Streptococcus mutans* forms consistent spatial arrangements in biofilm co-culture with other oral microbiota species. In this case, *S. mutans* consistently produce core clonal regions, around which form layers of non-*mutans* streptococci followed by non-streptococci. The metabolic activity of *S. mutans* within the inner regions of these multispecies biofilms caused low local pH that could recapitulate the rapid demineralization of enamel that occurs during development of caries *in vivo*.

Our work here highlights how the details of early surface colonization conditions can cascade into qualitative differences in subsequent biofilm architecture and ecological dynamics. This result points to several important directions for future work. Any phenotypes that alter surface exploration or settling patterns, including gliding and twitching motility as well as any positive or negative interactions within and between species – for example, via shared adhesin production, metabolite trophic interaction, or toxin secretion – could also cascade to major differences in biofilm spatial architecture. Differences in environmental topography and the orientation of nutrient supply, which may often derive in natural environments from the underlying surface rather surrounding liquid, should also be pursued to gain a fuller picture of how the subtleties of biofilm growth in realistic environments impact community structure. Novel experimental systems that implement these increases in ecological realism while maintaining access by high resolution imaging will be crucial platforms for further study.

## Materials and Methods

### Microfluidic assembly

Microfluidic chambers for biofilm growth were produced with polydimethylsiloxane (PDMS) using standard soft lithography techniques (103, 104). PDMS was cured on molds of chamber sets, which were then cut out, plumbed for inlet and outlet tubing, and bonded to #1.5 36mm by 60mm glass coverslips using plasma cleaning treatment. Biofilm growth chambers in this study measured 3000μm × 500μm × 75μm (L×W×D). Flow was manipulated using Harvard Apparatus Pico Plus syringe pumps loaded with 1ml BD plastic syringes. These syringes had 25-gauge needles affixed to them that were fitted with #30 Cole Palmer PTFE tubing with an inner diameter of 0.3mm. Tubing lines were placed into the inlet of each biofilm growth chamber, with outlet tubing feeding to effluent collection dishes.

### Bacterial strains and culture conditions

*V. cholerae* N16961 (O1 biovar El Tor) and *E. coli* AR3110 (a derivative of the K-12 strain W3110 with its cellulose operon restored (105)) were all grown overnight in lysogeny broth (LB) in shaking culture conditions at 37°C. *B. bacteriovorus* 109J stocks were obtained via co-culture with prey and filtering as described previously (35). For *V. cholerae*, constitutive fluorescent protein expression constructs, gene deletions, and -FLAG fusions to native *rbmA*, *rbmC*, and *bap1* loci were introduced onto the chromosome using standard allelic exchange methods (25, 35). For *E. coli*, fluorescent protein expression constructs were introduced in single copy on the chromosome using Lambda-red recombination (106). For *B. bacteriovorus*, GFP expression was driven by plasmid borne expression *in trans* (35).

### Biofilm growth and matrix immunostaining

To introduce bacteria to the microfluidic devices, overnight cultures of *V. cholerae* and *E. coli* were diluted to OD_600_ = 1.0 and inoculated into chambers via micropipette. The inoculating cultures were left to stand in the chambers for 1 h without flow to allow for surface colonization. Flow of M9 minimal media with 0.5% glucose was then initiated at 0.2μL/min. For dual species prey biofilms with *V. cholerae* and *E. coli* inoculated together, cultures of each species were each diluted to OD_600_ = 1.0 and mixed 1:1 just prior to inoculation into chambers. For experiments involving matrix staining, biofilms inoculated with *V. cholerae* strains producing FLAG-labeled matrix proteins RbmA, RbmC, or Bap1 were grown as above until 48 h, at which point the inflow syringes were changed to new syringes containing M9 minimal media with 0.5% glucose and Cy3-conjugated anti-FLAG antibody at 1μg/mL, maintaining the same flow rate of 0.2μL/min. After an additional 4 h, biofilms were imaged by confocal microscopy as detailed below.

### Bacterial predator introduction and invading bacteria introduction

To introduce bacterial predators, *B bacteriovorus* were diluted to OD_600_ = 1.0 (2.5 × 10^9^ PFU/mL) using M9 with 0.5% glucose. These cultures were then loaded into 1ml syringes. At the time of predator inoculation, the syringe and tubing containing sterile media were changed for the syringe and tubing containing predators, and flow was resumed at 0.2μL/min. After 1 h, the tubing was changed back to an influx of sterile M9 with 0.5% glucose. For the introduction of invading *E. coli* (SI Figure S8), cultures of *E. coli* were concentrated to OD_600_ = 5.0 prior to inoculation and loaded into 1 mL syringes. A similar tubing swap was performed as described above and *E. coli* flow was allowed to continue for 24 h. After 24 h, the syringe and tubing were replaced with media containing predators as described above. After 1 h of predator introduction, the syringe and tubing containing sterile M9 media with 0.5% glucose were replaced once again to resume sterile medium influx.

### Bacterial dispersal and re-colonization to determine *E. coli* viability after predation

Co-culture biofilms were grown in standard conditions for 48 h as described above and then imaged prior to introduction of *B. bacteriovorus*. Following this first imaging step, *B. bacteriovorus* was introduced as described above; 48 h later, biofilms were imaged again. Biofilms were then dispersed by removing the tubing from the microfluidic devices before vigorously pipetting M9 media and air bubbles back and forth between the inlet and outlet ports of the biofilm chambers; this technique most effectively removes the majority of the *E. coli* population residing within highly packed *V. cholerae* microcolonies (78). The resulting cell suspension was vortexed before being serially diluted and plated for *E. coli* CFUs. CFUs were determined by plating onto LB agar with 50μg/mL Kanamycin to distinguish the two prey species; *E. coli* strain carried a kanamycin resistance cassette along with its constitutive fluorescent protein expression construct, while *V. cholerae* was not Kanamycin resistant. In a separate experiment, the cell suspension collected was used to inoculate new microfluidic devices, which were imaged 24 h after initial colonization to assess subsequent *E. coli* growth.

### *E. coli* biofilm growth with *V. cholerae* supernatants containing labeled matrix proteins

*V. cholerae* matrix supernatants were prepared by growing strains harboring a clean deletion of *vpsL*, without which *V. cholerae* cannot produce the matrix polysaccharide VPS; loss of VPS production prevents the retention of the matrix proteins RbmA, RbmC, and Bap1, so that they are more readily released into the supernatant. A D*vpsL* strain with a -FLAG tag fused to the chromosomal locus *rbmA*, *rbmC*, or *bap1* was used to produce RbmA-FLAG, RbmC-FLAG, or Bap1-FLAG supernatants, respectively. To prepare the supernatants, each of these strains was grown overnight, centrifuged to pellet bacterial cells, and the supernatants was removed by micropipette and passed through a 0.22 μm filter to ensure removal of all *V. cholerae* cells.

To prepare *E. coli* biofilms, overnight cultures of *E. coli* were inoculated into microfluidic devices for biofilm growth as described above. At 48 h after the start of biofilm growth, in three separate treatments (one for each matrix protein), the prepared supernatants containing RbmA-FLAG, RbmC-FLAG, or Bap1-FLAG were mixed 1:5 into fresh M9 with 0.5% glucose and 1% BSA and loaded into syringes that were swapped to the inlets for the *E. coli* monoculture biofilms. At 72 h, the media syringes leading into the chambers were replaced again with new syringes containing fresh M9 minimal media with 0.5% glucose and 0.1 μg/ml Cy3-conjugated anti-FLAG antibody. Chambers were then incubated with the new input media overnight. Biofilms were then imaged by confocal microscopy to assess the association of strained matrix protein from the supernatants with the *E. coli* monoculture biofilms.

### Microscopy and image analysis

Imaging was performed on Zeiss LSM 880 and LSM 980 laser scanning confocal microscopes. Both instruments were fitted with a 40x/ 1.2 N.A. water objective and a 10x/ 0.4 N.A. water objective. 488nm, 543nm, and 594nm laser lines were used to excite the fluorescent proteins GFP (*B. bacteriovorus*), mKO-k (*E. coli*), and mKate2 (*V. cholerae*), respectively. For experiments that included Cy3-labeled matrix proteins, the 543nm laser line was used to excite Cy3. In these experiments requiring Cy3 imaging, we used *E. coli* producing the fluorescent protein mTFP1 (excited with a 488nm laser) to ensure no overlap between fluorescent proteins used for bacterial identification and Cy3 emission spectra. The hardware was controlled by ZEN Black for the LSM 880 and ZEN blue for the LSM 980. To obtain data for image analysis, 3 or more image stacks were taken within each chamber and averaged to produce each biological replicate in independent chambers. Prior to export, the emission channels were processed by linear unmixing and constrained iterative deconvolution (Poisson likelihood, Zero order regularization, Newton Raphson optimization, maximum 40 iteration, 0.1% quality threshold, scalar theory PSF generation) in ZEN blue. These images were then exported and analyzed using BiofilmQ. 3-D image renderings were produced using Zen or VTK export from BiofilmQ and subsequent processing in ParaView.

### Quantification and statistics

Biofilms were segmented using the image analysis framework BiofilmQ (107). Briefly, BiofilmQ denoises image data and segments biovolume into pseudo-cell cubes that can be used for spatially resolved data analysis. Image files were imported to BiofilmQ directly from ZEN. For population level experiments, total biovolume of each species was tracked. For data involving micro-scale architecture, only 40x data were used and were segmented into cubes with a volume of 2.06μm^3^. Cell packing was determined using a local density calculation with a spatial range of 6μm. This is the spatial scale – neighborhood volume fraction – that we have previously shown to be most relevant pertinent to measuring architectural packing-based protection against *B. bacteriovorus* (35). The degree of predation was obtained by calculating Manders overlap coefficients between the prey species and *B. bacteriovorus*. If any amount of predator signal is detected within prey signal, the degree of predation will be above 0. Mann-Whitney U-tests with Bonferroni correction were used for pairwise comparisons.

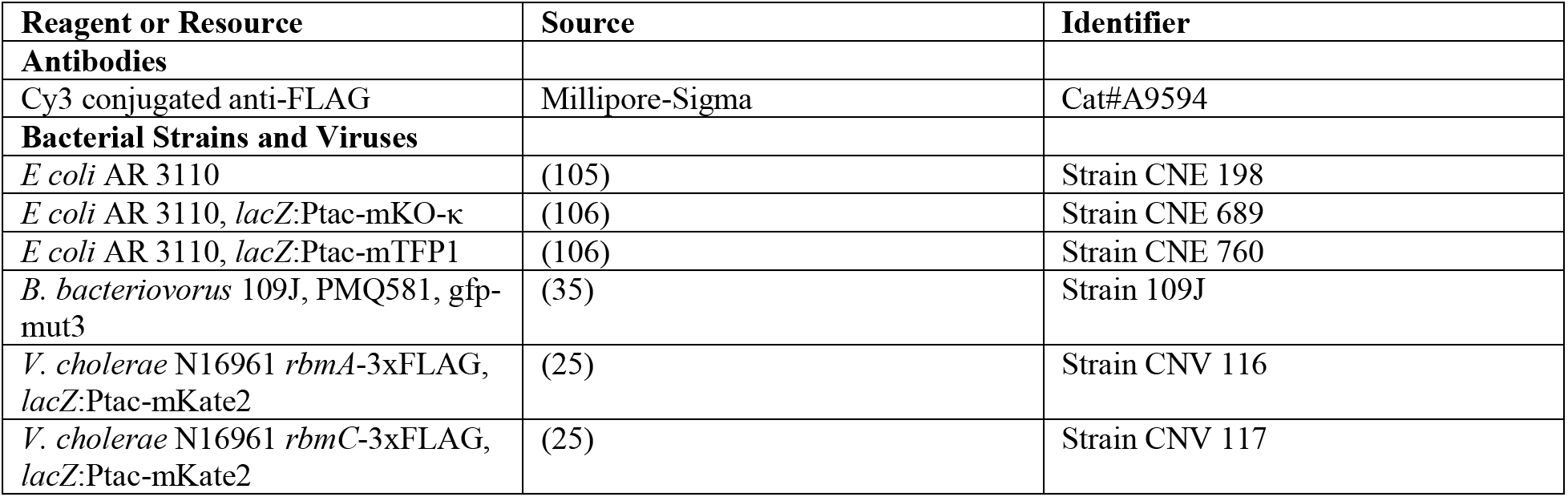
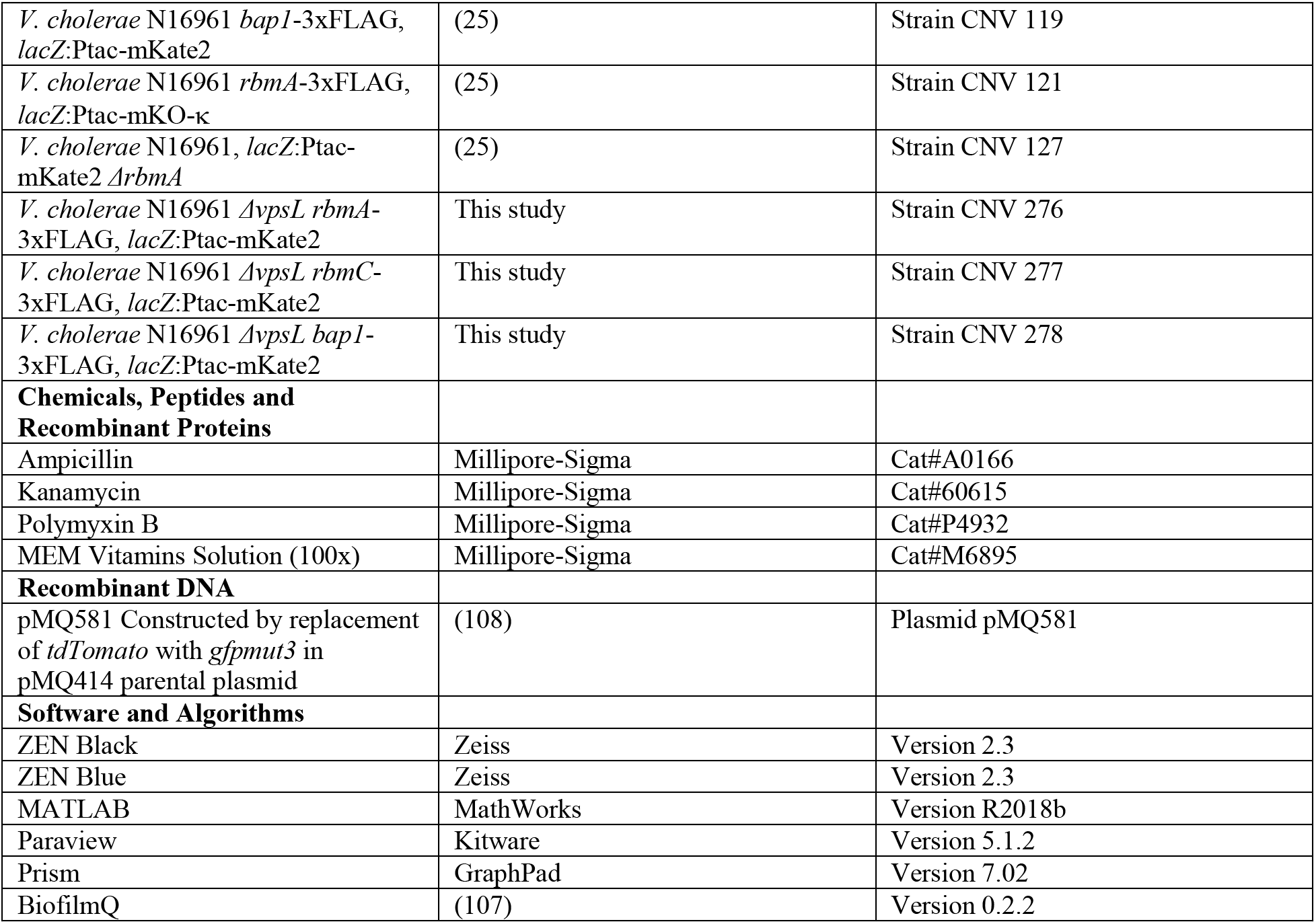

## Author Contributions

BRW and CDN conceived the study. BRW, JBW, and CDN designed experiments. CDN acquired funding for the work and supervised the project. ME and DEK provided reagents/tools. BRW and JBW performed experiments, image processing of microscopy data, and figure drafting. BRW and CDN analyzed data, finalized the figures, and wrote the paper.

## Classification

Biological Sciences: Microbiology

## Acknowledgements

We are grateful to Blair Costelloe, Robert Cramer, Matthew Ayres, and Jing Yan for helpful advice on earlier versions of the manuscript, and to members of the Nadell Lab at Dartmouth for their input during development of the experiments and analysis. BRW received support from a Gillman Fellowship from the Department of Biological Sciences at Dartmouth. JBW was supported by a GAANN Fellowship from the Dartmouth Department of Biological Sciences. CDN received support from the Simons Foundation award number 826672, NSF grant IOS 2017879, NSF grant MCB 1817352, and grant RGY0077/2020 from the Human Frontier Science Program.

## Supporting Information

**Figure S1.**
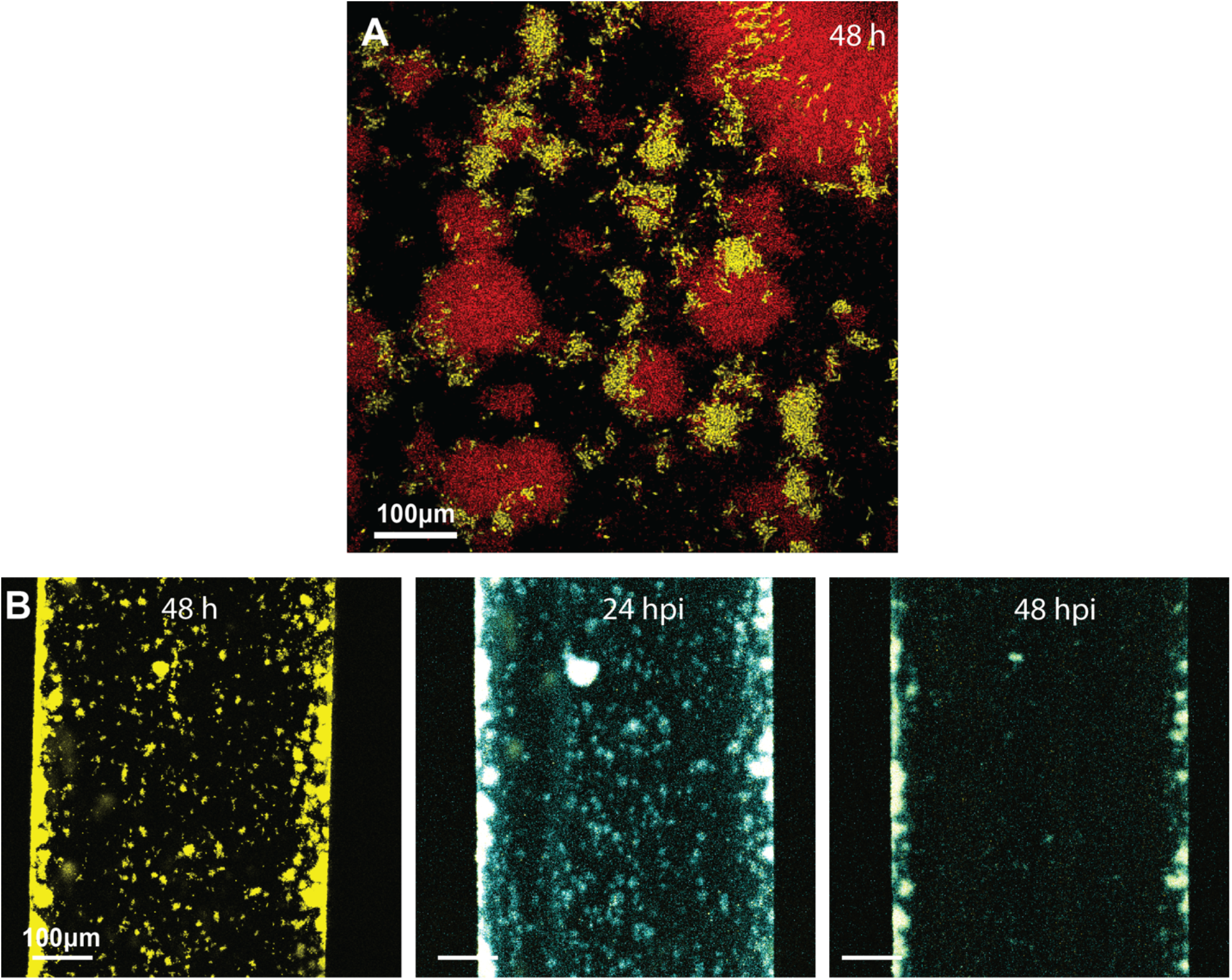
(A) A representative image of cohabitation of *E. coli* (yellow) and *V. cholerae* (red) and variable colony structure in microfluidic culture conditions after 48 h of growth following surface colonization. (B) *E. coli* mono-culture biofilm representative images at 48 h of growth, as well as 24 and 48 hours post introduction (hpi) of predator *B. bacteriovorus* (cyan). The few remaining *E. coli* cells present at 48 hpi can be seen still undergoing predation by *B. bacteriovorus* and ultimately die as well. All images here are single optical sections taken just above the glass substratum on which biofilms are growing.

**Figure S2.**
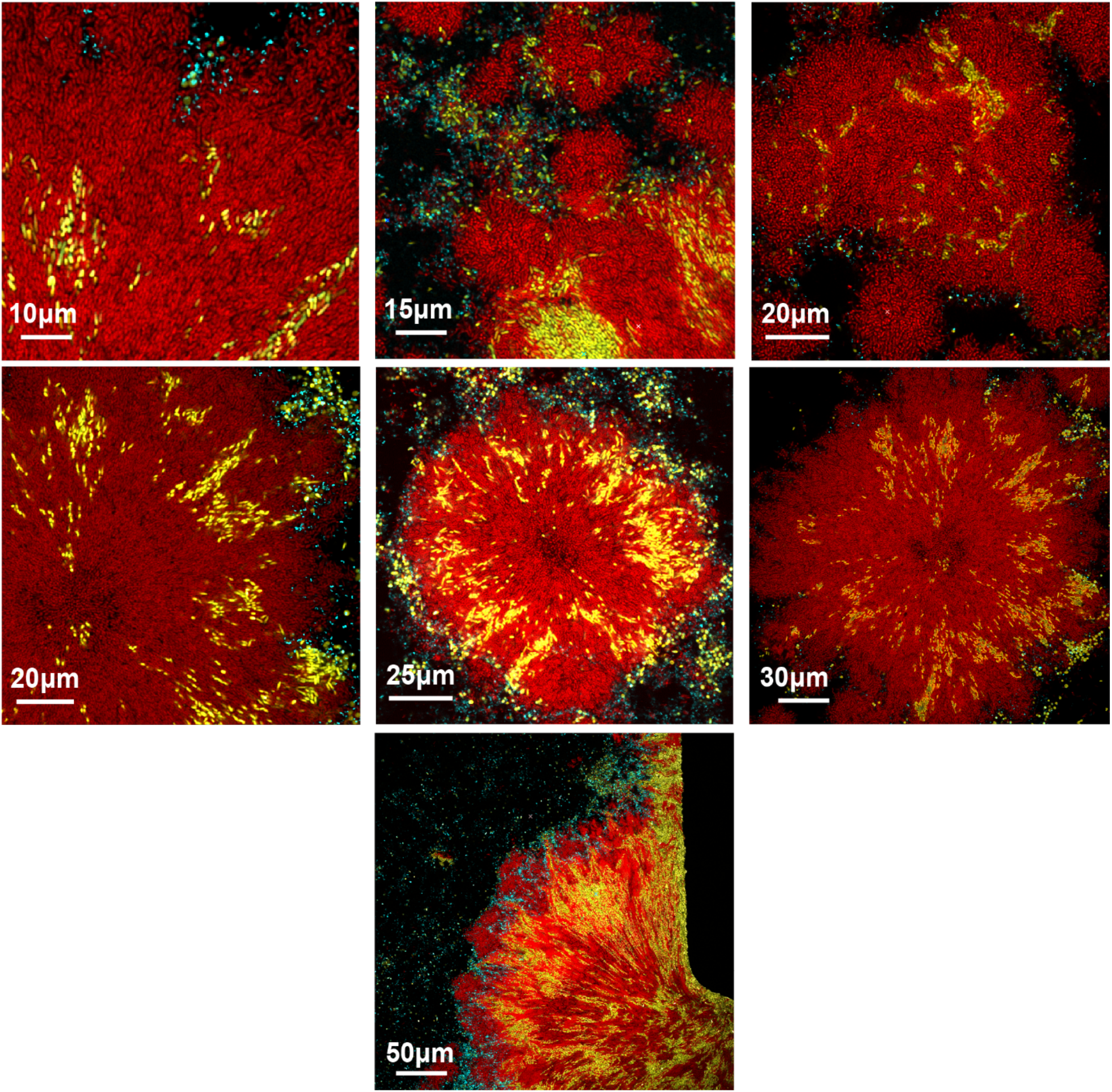
Representative examples of *E. coli* (yellow) inclusions within conventional *V. cholerae* (red) biofilm structures with high cell packing; introduced *B. bacteriovorus* can be seen in cyan. All of the images here are single x-y optical sections taken just above the glass substratum on which the biofilms are growing. The amount and size of *E. coli* clusters within each *V. cholerae* cell group varies from one instance to another, but in all cases *E. coli* cell groups enveloped within *V. cholerae* colonies in this manner are restricted to the bottom layers of the growing biofilm, just above the glass.

**Figure S3.**
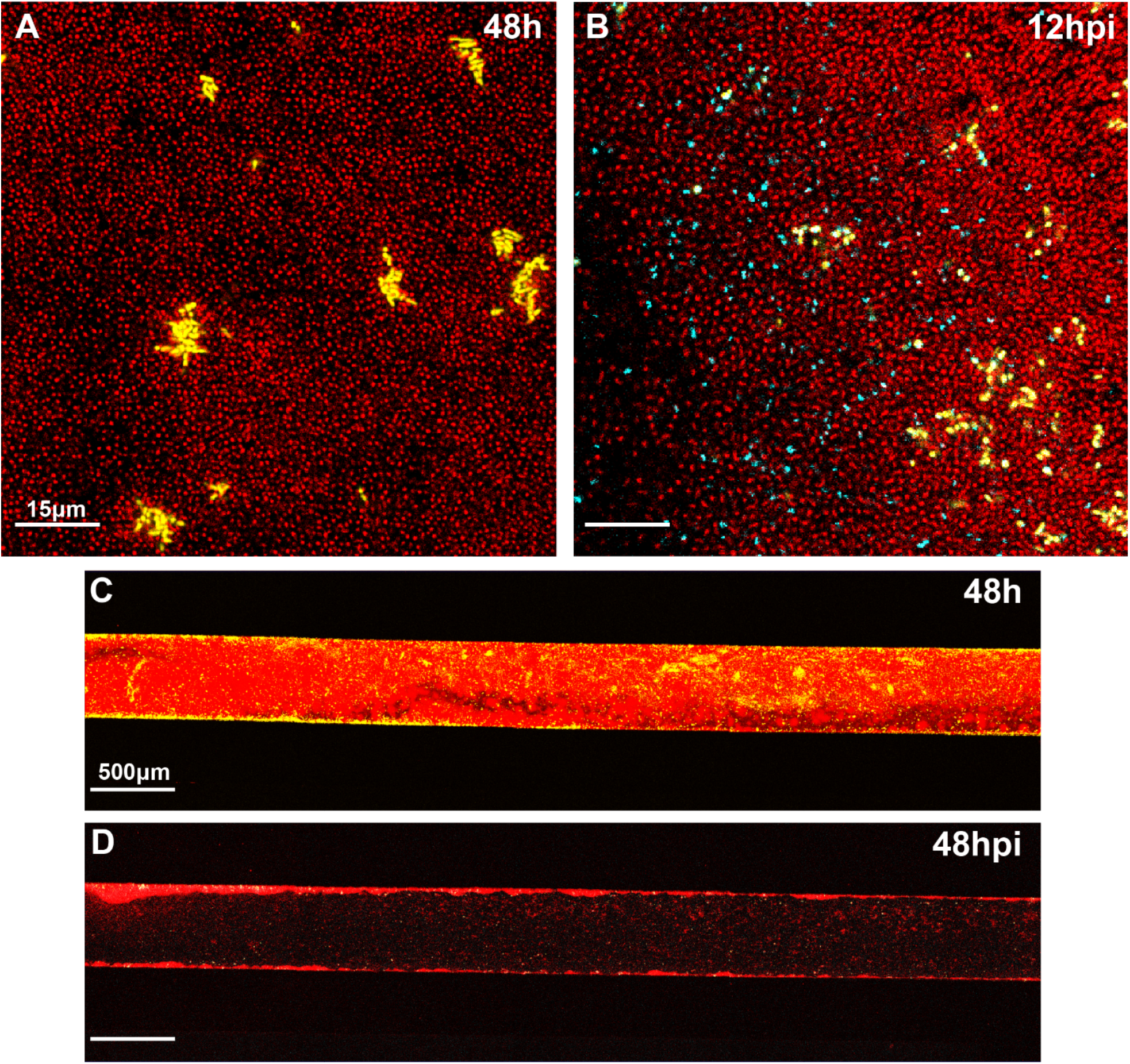
*ΔrbmA V. cholerae* (red) biofilms cannot protect *E. coli* (yellow) inclusions from predation by *B. bacteriovorus* (cyan). *E. coli* and *ΔrbmA V. cholerae* biofilm growth at (A) 48 h and (B) 48 h post introduction of predators (hpi). (C-D) Whole microfluidic chamber images showing *E. coli* and *ΔrbmA V. cholerae* biofilm growth (C) before and (D) after predation.

**Figure S4.**
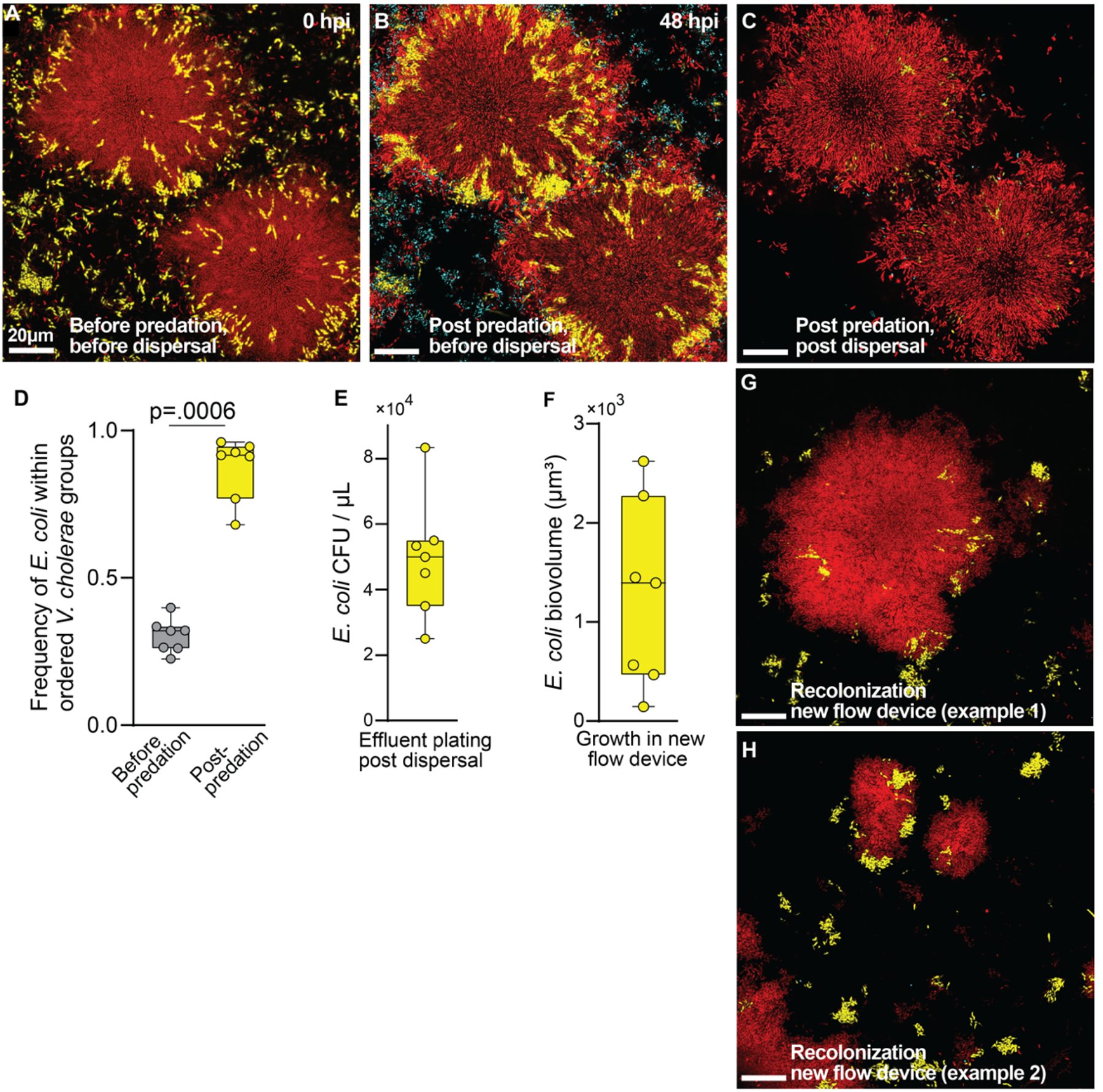
Dispersal of dual species biofilms post predation shows that *E. coli* (yellow) growing within mature *V. cholerae* (red) colonies remain viable and can colonize and grow in downstream locations. (A) Image of *V. cholerae* and *E. coli* in biofilm co-culture prior to introduction of predatory *B. bacteriovorus* (cyan). (B) Subsequent image of same location as in (A), 48 h post introduction of predatory *B. bacteriovorus*, which can be seen consuming loosely packed cells outside of the two larger central *V. cholerae* biofilm clusters. Both of the high packing clusters have enveloped pockets of *E. coli* embedded in them that are also protected from *B. bacteriovorus* exposure. (C) After predators have killed exposed cells, biofilms are subjected to a disturbance event that disperses most of the *E. coli* from highly packed clusters. (D) Fraction of viable *E. coli* (i.e., without predators inside them) within the boundaries of highly packed *V. cholerae* cells groups before predation and after predation (*n*=7). (E) Colony forming unit quantification of viable *E. coli* in liquid effluent following dispersal. (F) Quantification of *E. coli* biofilm production in new sterile chambers colonized by the effluent produced from the dispersal effluent (images taken 24 h after colonization of the new sterile chambers). (G,H) Representative image examples of the early dual species biofilms produced by *V. cholerae* and *E. coli* in fresh chambers colonized by the effluent from the dispersed chambers (example in panel C).

**Figure S5.**
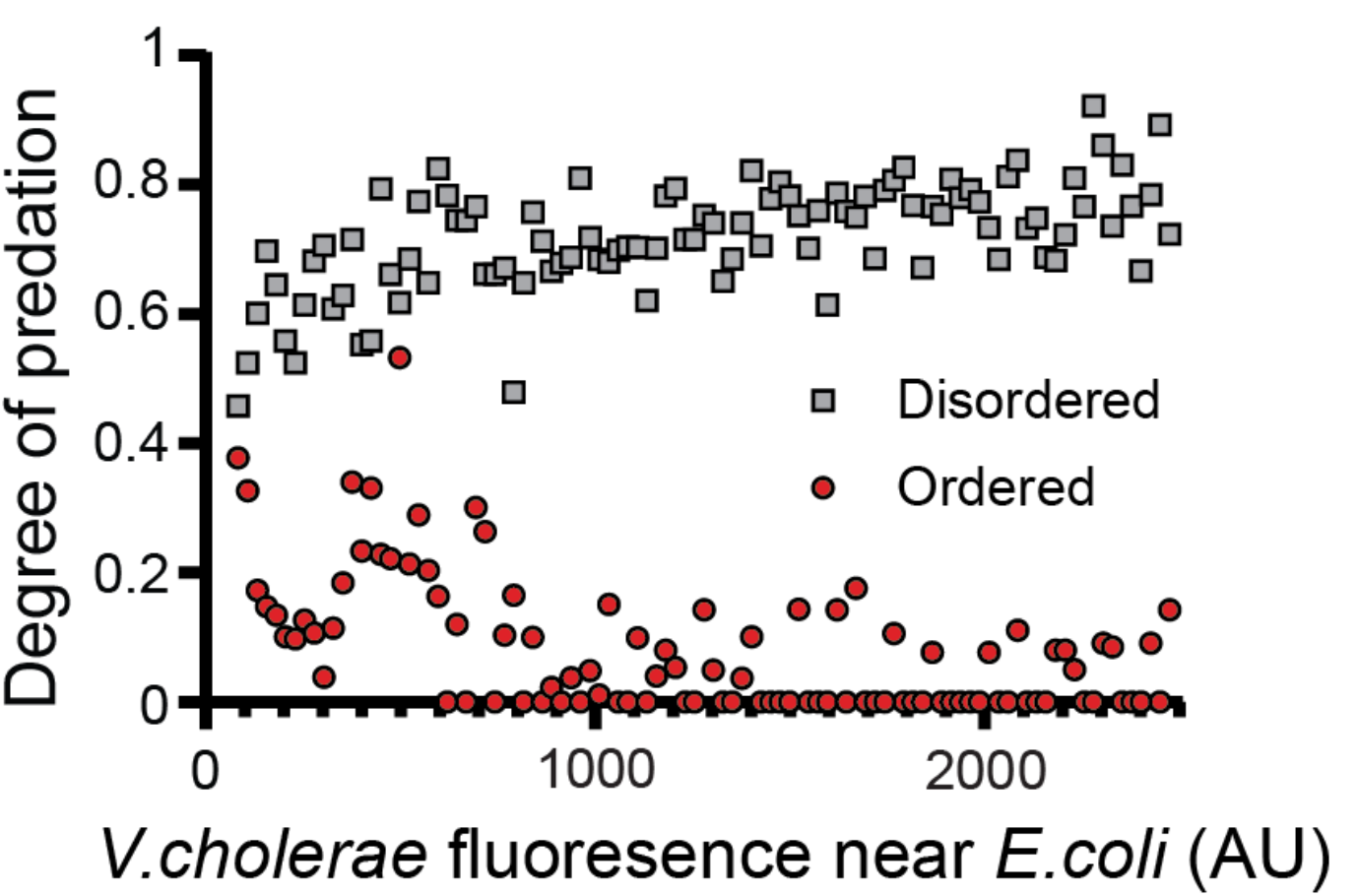
Whether *E. coli* is protected from predation is determined by the colony type in which it is embedded, rather than simply a function of *V. cholerae* fluorescence in its immediate vicinity. The inverse is also true for *V. cholerae,* as shown in the main text Figure 3E.

**Figure S6.**
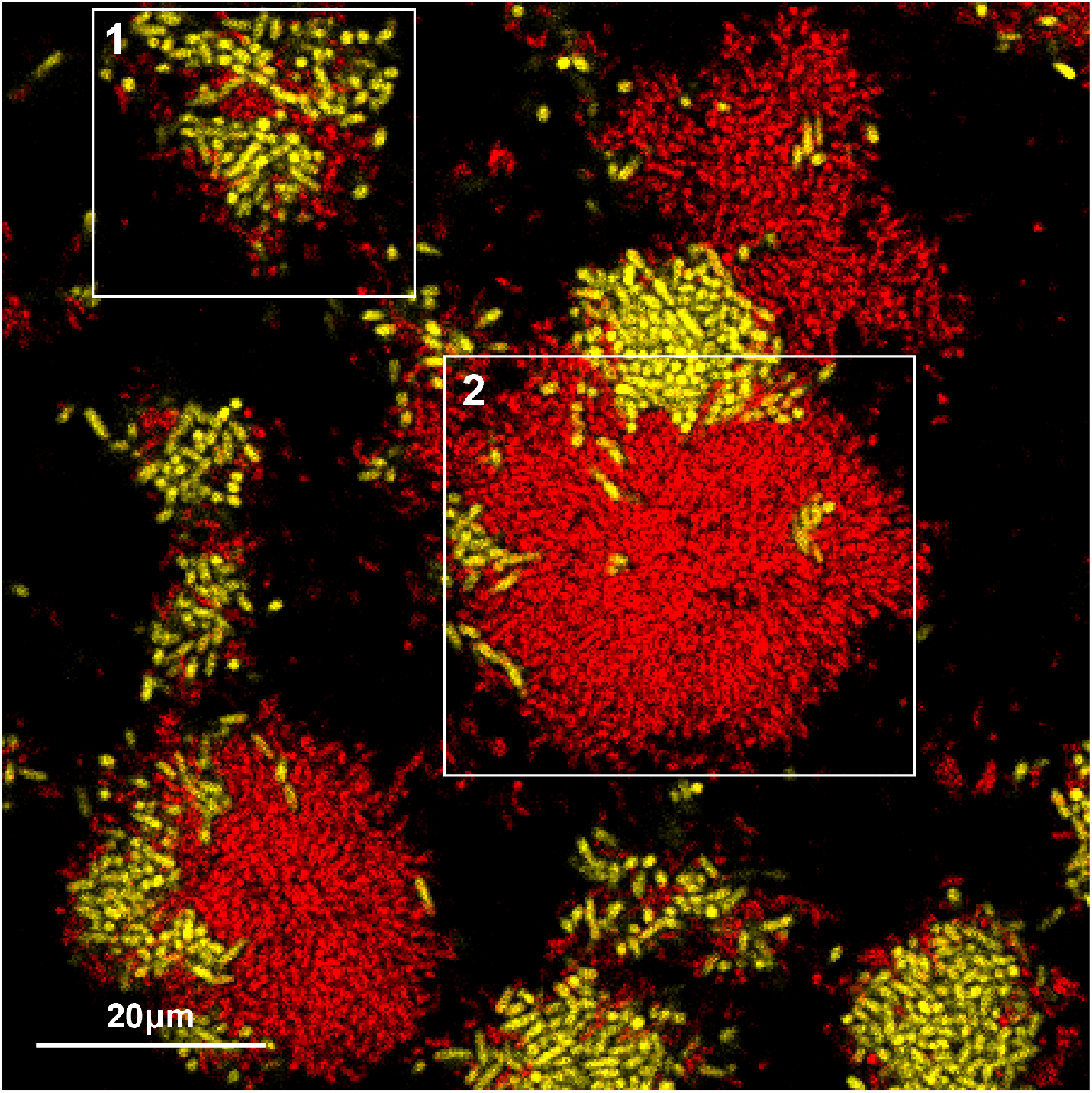
A supplemental representative image of variable multispecies microcolony architecture in biofilms of *V. cholerae* (red) and *E. coli* (yellow). The two qualitatively different colony structures we observe (ordered vs. disordered) can both occur in close proximity (1: disordered; 2: Ordered)

**Figure S7.**
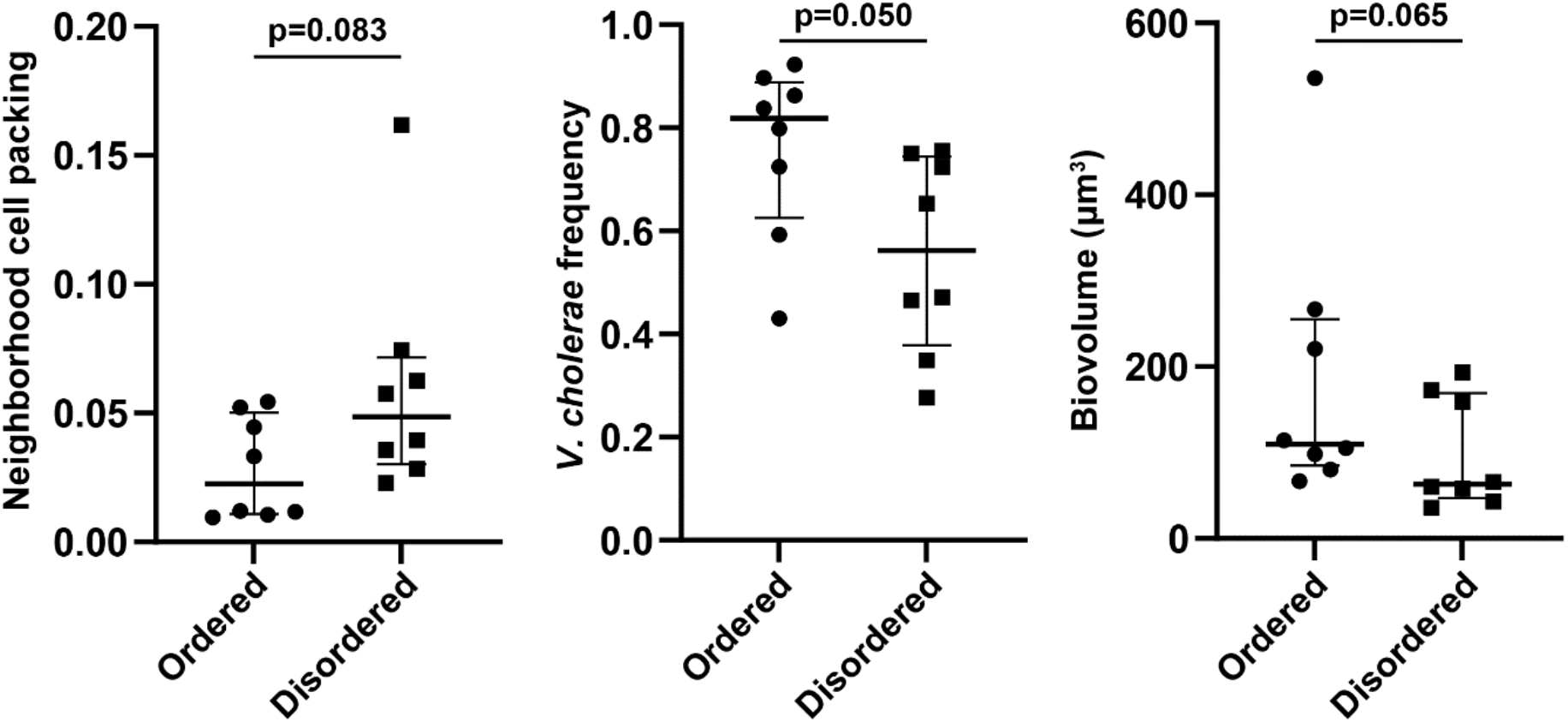
Relative to initial distance between *V. cholerae* and *E. coli* cells (see main text Figure 4G), measures of cell packing, *V. cholerae* relative abundance, and total biovolume at the start of biofilm growth were not significantly different between conditions that would ultimately produce ordered high-packing microcolonies versus disordered microcolonies.

**Figure S8.**
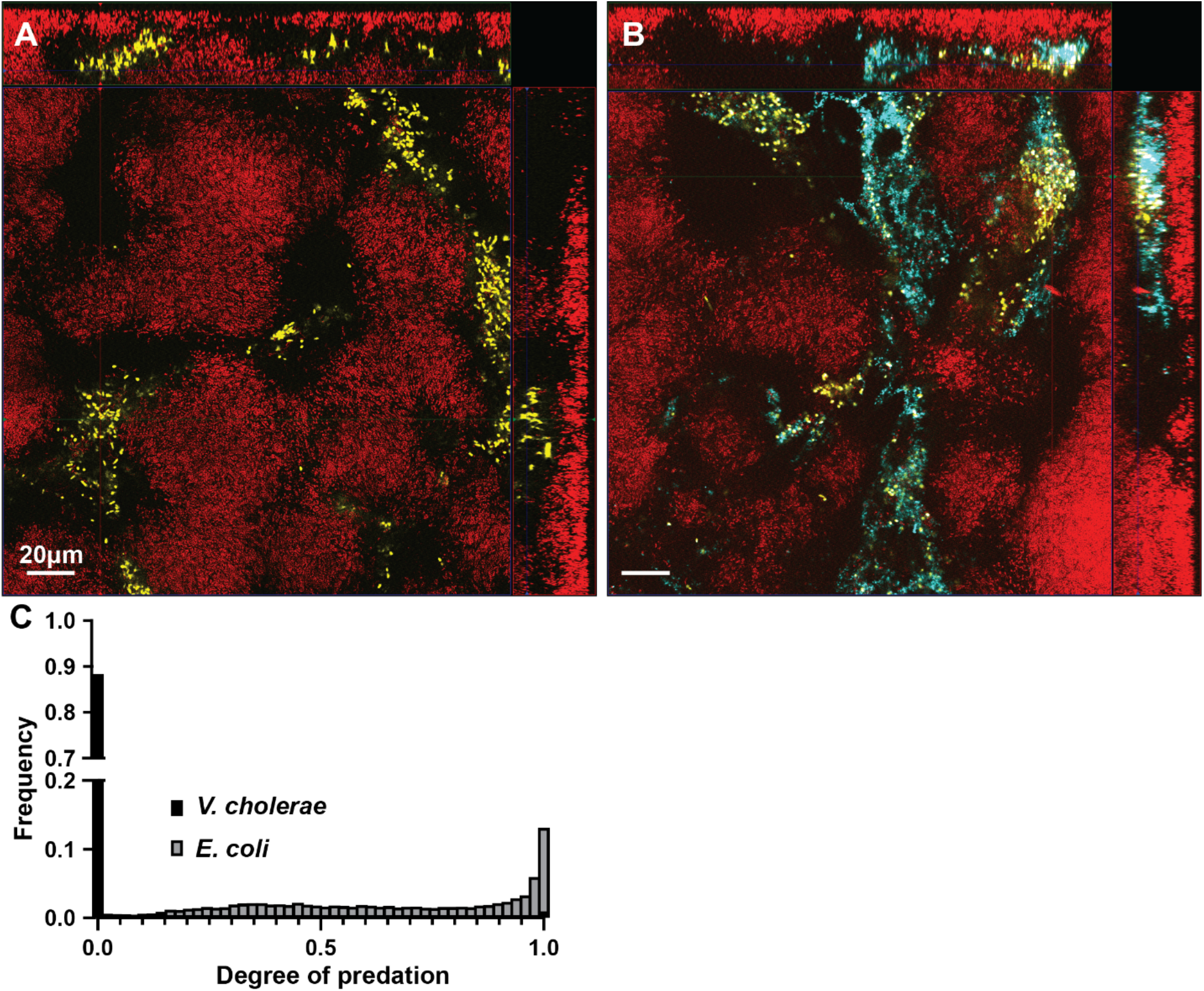
Continuous introduction of *E. coli* (yellow) after *V. cholerae* (red) biofilm establishment does not change the morphology of *V. cholerae* biofilms. (A,B) Representative images of *V. cholerae* biofilms grown for 48 h prior to the continuous introduction of *E. coli* cells in the media: (A) before and (B)1 h after introduction of *B. bacteriovorus* (cyan). *E. coli* is not able to invade or disrupt the structure of pre-existing highly packed *V. cholerae* biofilms. (C) Frequency distribution of predation for *V. cholerae* and *E. coli* in this experiment.

**Figure S9.**
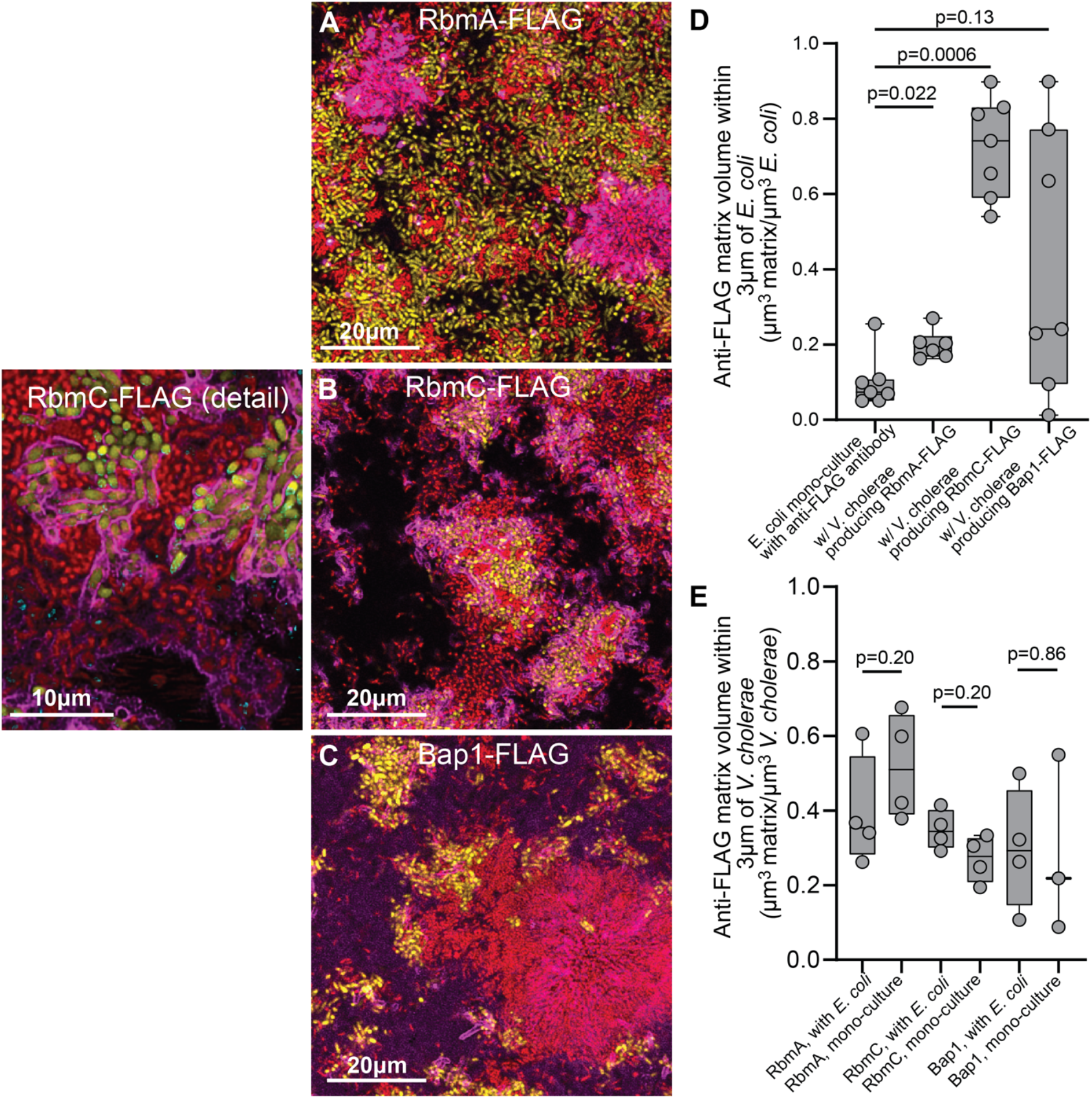
Localization of *V. cholerae* secreted matrix proteins (magenta) RbmA, RbmC, and Bap1 in coculture biofilms of *V. cholerae* (red) and *E. coli* (yellow). (A-C) Representative images of co-culture biofilms of *E. coli* and *V. cholerae* producing -FLAG labeled versions of (A) RbmA, (B) RbmC (with higher magnification at left for detail to show localization of RbmC to *E. coli* exterior), and (C) Bap1-FLAG. FLAG-tagged matrix proteins were stained with monoclonal anti-FLAG antibody conjugated to Cy3. (D) Quantification of segmented secreted matrix volume in proximity to *E. coli* (normalized to volume of *E. coli*), compared with an antibody control (left-most bar) in which monoculture *E. coli* biofilms were grown with addition of only the anti-FLAG antibody to assess background staining. Pairwise comparisons denote Mann-Whitney U tests with *n* = 7. After Bonferroni correction, only the pairwise comparison of RbmC localization to *E. coli* and the antibody control in panel D is statistically significant at p<0.05. (E) Comparison of matrix protein localization in proximity to *V. cholerae* (normalized to volume of *V. cholerae*) in monoculture controls relative to biofilms in co-culture with *E. coli*. Pairwise comparisons denote Mann-Whitney U tests with *n* = 3-4.

**Figure S10.**
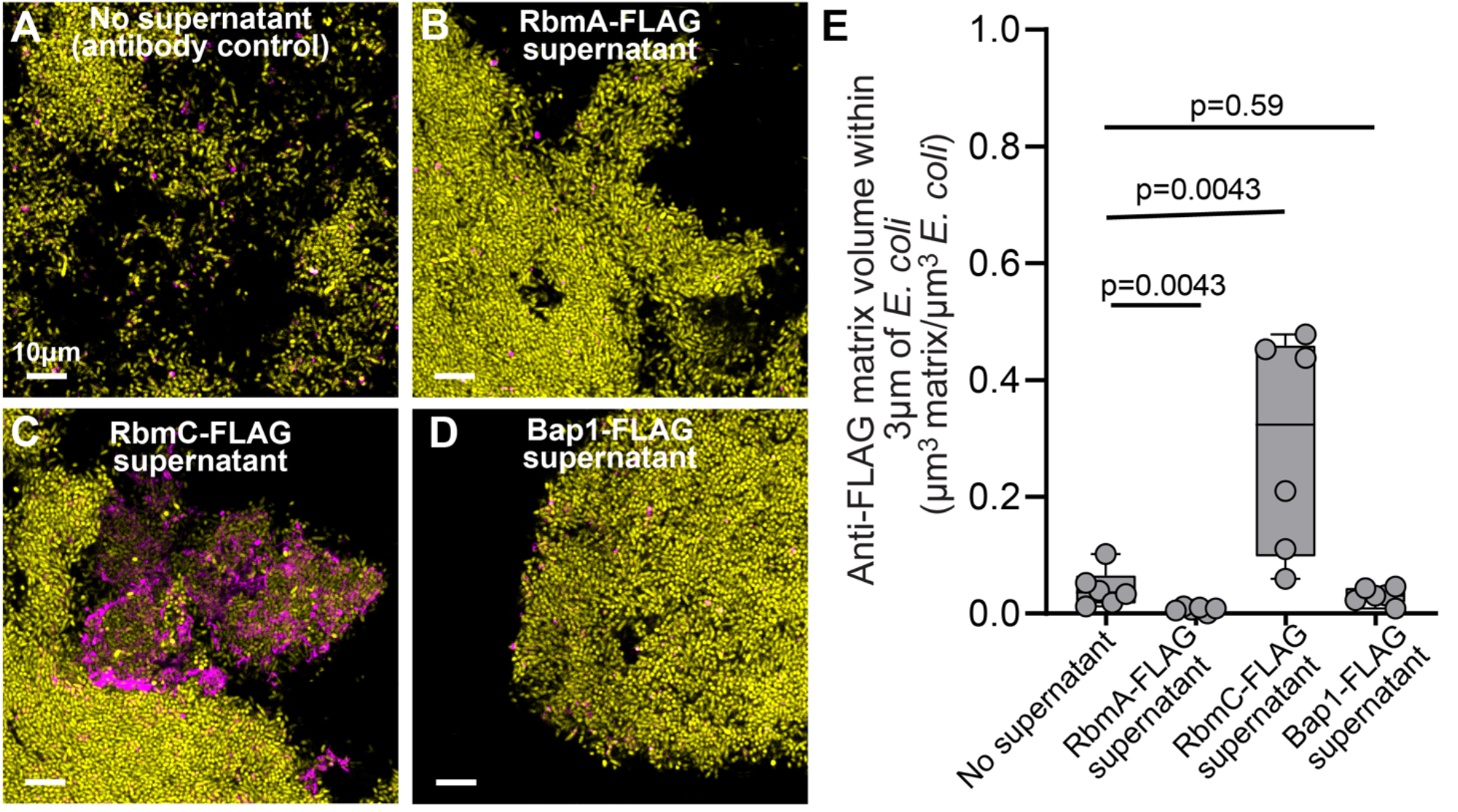
Localization of FLAG-labeled matrix proteins RbmA, RbmC, and Bap1 from cell-free supernatants of *V. cholerae* added to monoculture biofilms of *E. coli*. (A-D) Representative images of *E. coli* (yellow) monoculture biofilms following addition of supernatants containing (A) Anti-FLAG antibody only, (B) RbmA-FLAG with anti-FLAG antibody, (C) RbmC with anti-FLAG antibody, and (D) Bap1 with anti-FLAG antibody. Anti-FLAG Cy3 fluorescence is shown in magenta. (E) Quantification of stained matrix volume in proximity to *E. coli*, normalized to volume of *E. coli*, from the supernatant experiments depicted in representative images A-D. As for the co-culture experiments shown in Figure S9, RbmC-FLAG accumulates around *E. coli* cells significantly above background.

**Figure S11.**
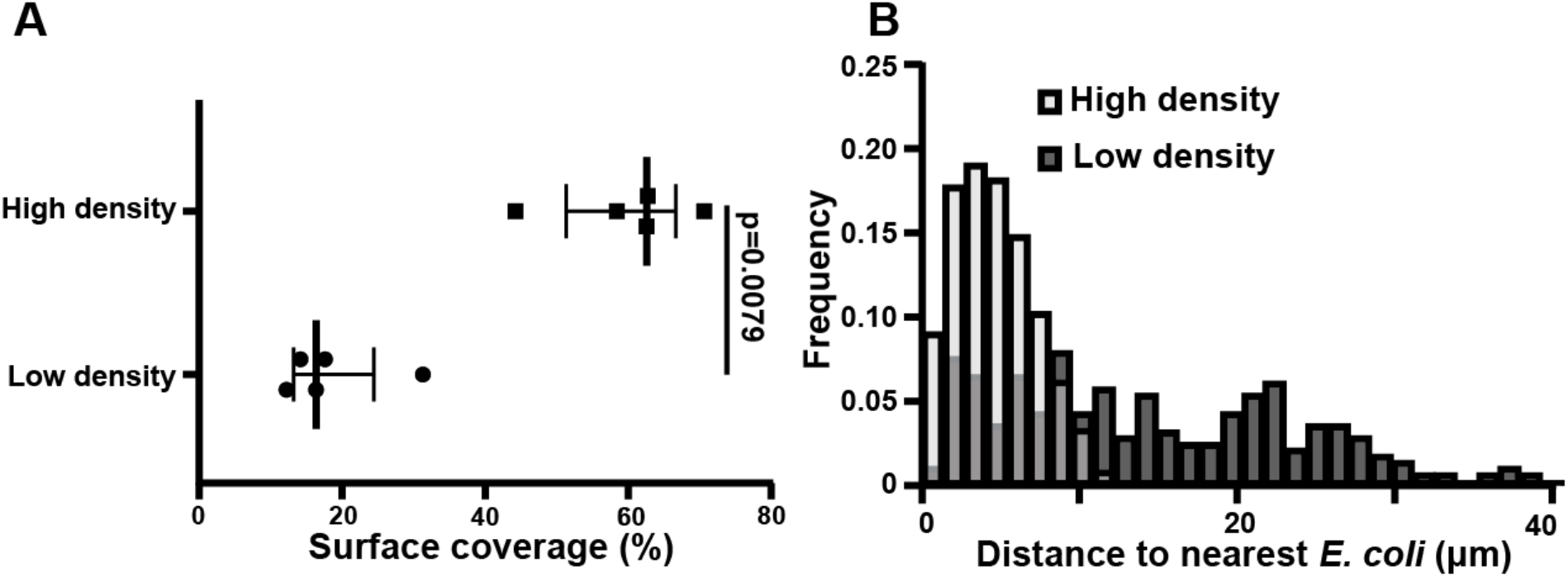
Surface coverage and distributions of distances between *V. cholerae* and *E. coli* in high versus low surface colonization density experiments in Figure 5 of the main text. (A) Comparison of average surface colonization between high-and low-density colonization conditions. Pairwise comparison denote Mann-Whitney U-tests (*n =* 5). (B) Frequency distributions of the average distance to the nearest *E. coli* cell for all *V. cholerae* in low versus high surface colonization density settings.

